# Accurate prediction of nitrogen fixation in cyanobacteria reveals the dynamic evolution driving high retention rate with mosaic distribution

**DOI:** 10.64898/2026.01.15.699626

**Authors:** Kazuma Uesaka, Yuichi Fujita

## Abstract

Cyanobacteria are the only prokaryotic group capable of simultaneously performing oxygenic photosynthesis and nitrogen fixation by nitrogenase extremely sensitive to oxygen. However, systematic understanding of the phylogenetic distribution of nitrogen-fixing ability remains incomplete because of difficulties in accurately identifying genes based on genome information due to occasional confusion with nitrogenase-like enzymes and the high degree of fragmentation in *nif* genes in heterocystous species. In this study, we comprehensively evaluated the distribution of the minimal nitrogen fixation gene set (*nifHDKENB*) across 586 major cyanobacterial strains using a 2-dimensional similarity plot with the negative logarithm of the e-values and protein length allowing accurate prediction of nitrogen fixation. The evaluation that almost all heterocystous strains are nitrogen fixer supports the accuracy of this method. Overall, 48% of the strains were predicted to be nitrogen-fixing with a mosaic distribution in non-heterocystous strains. Using the results as training data, nitrogen fixation was evaluated for a broader cyanobacterial lineage, including uncultured strains and metagenomes. Approximately 20% of 2,718 strains were evaluated to be nitrogen fixers, revealing a higher retention rate in Cyanobacteria phylum than other bacterial phyla. We also found a novel *nif* genes belonging to Group II and discussed the dynamic evolution process in cyanobacteria.

## Introduction

Nitrogen fixation is the process of converting atmospheric nitrogen (N_2_) into ammonia (NH_3_) that is available to most organisms. Biological nitrogen fixation, catalyzed by the metalloenzyme nitrogenase, plays a critical role in the global nitrogen cycle. Identifying nitrogen-fixing organisms and determining their distribution and abundance are essential for understanding the nitrogen cycle on a global scale [1,2]. Nitrogenase is distributed across 16 phyla in Bacteria and 1 phylum in Archaea, however, it occurs a patchy manner within each phylum [3,4].

Nitrogenase consists of two easily separable protein components: the Fe protein and the MoFe protein. The Fe protein is a homodimer of the NifH protein containing a single [4Fe-4S] cluster. The Fe protein is reduced by ferredoxin, and transfers electrons to the catalytic component, the MoFe protein, coupled with ATP hydrolysis. The MoFe protein is a heterotetramer composed of two subunits, NifD and NifK, which show low sequence similarity. The MoFe protein contains two pairs of metal centers: the P-cluster ([8Fe-7S]) and the iron-molybdenum cofactor (FeMo-co, [1Mo-7Fe-9S-C-homocitrate]). Electrons from the Fe protein is transferred via the P-cluster to FeMo-co, where the FeMo-co bound N_2_ molecule is reduced. FeMo-co is synthesized by the actions of NifB and NifEN. NifB synthesize the precursor of FeMo-co, and NifEN acts as a scaffold for FeMo-co assembly. The minimal unit to produce functional nitrogenase is proposed to be the six genes: the structural genes, *nifHDK*, and *nifENB* involved in FeMo-co biosynthesis.

The three metal clusters of nitrogenase and their precursors are extremely vulnerable to oxygen. Nitrogenase undergoes irreversible degradation within seconds or minutes after exposure to oxygen [5]. Thus, nitrogen fixation requires highly anaerobic environment. Nitrogen-fixing organisms inhabiting aerobic or microoxic environments have developed diverse mechanisms to protect nitrogenase from oxygen [6,7]. While the nitrogenase is highly conserved even across different taxa [8], the protection systems are significantly diverse among species.

Cyanobacteria are prokaryotes performing oxygenic photosynthesis and represent the only group of organism capable of coexisting both oxygenic photosynthesis and oxygen-sensitive nitrogen fixation. They employ distinct strategies to solve the Oxygen Paradox. Most known strategy is the spatial separation of nitrogen fixation and photosynthesis through heterocyst differentiation in filamentous species. In the well-studied species *Anabaena* sp. PCC 7120, at a rate of approximately 1 in every 8–15 vegetative cells along the filament differentiates into a heterocyst [9], a cell specialized for nitrogen fixation [10–12]. Heterocysts maintain an anaerobic cellular environment through high activity of respiration, production of thick cell walls, and dismantle of photosystem II [13–15]. This mechanism enables nitrogen fixation even under aerobic environments. In contrast, some unicellular nitrogen-fixing species employ the temporal separation, restricting nitrogen fixation to the night and photosynthesis to the day under circadian control [16]. However, other diverse strategies occurs in other strains: some unicellular cyanobacteria in microbial mats exhibit nitrogen-fixing activity only at dawn and dusk [17], while the non-heterocystous filamentous genus *Trichodesmium* performs nitrogen fixation during the day [18].

Nitrogen-fixing cyanobacteria play a crucial ecological role by providing nitrogen sources, particularly in oligotrophic environments [19]. Therefore, quantifying cyanobacterial nitrogen fixation is essential for understanding the global nitrogen cycle [20]. With advancements in sequencing technologies and metagenomic analyses, today approximately 8,200 cyanobacterial genomes are publicly available (December 2025) [21]. Leveraging this vast genomic dataset, it is expected to clarify the overall distribution of diazotrophic cyanobacteria based on genomic information [12].

Cyanobacteria are metabolically and morphologically diverse, making their evolutionary relationships challenging to classify. The morphology based classification system proposed in 1979 has been widely utilized [22], in which 178 strains were categorized into five groups: unicellular (Section I), unicellular forms with multiple fission (baeocytes; Section II), non-heterocystous filamentous (Section III), non-branching heterocystous filamentous (Section IV), and branching heterocystous filamentous (Section V). While strains in Sections IV and V are easily inferred to be nitrogen fixers due to the definition of heterocystous [23], evaluating the nitrogen-fixing potential for strains in Sections I–III is more challenging [24]. Out of 118 strains in Sections I–III tested for nitrogen fixation using acetylene reduction assay only 46 strains showed the activity [22]. The uneven distribution across Sections I–III suggested that nitrogen-fixing ability might not align with phylogenetic classification. Based on these and subsequent experimental results, it is estimated that approximately half of all cyanobacterial species possess nitrogen-fixing potential [7,25]. Although new molecular phylogenetic classifications based on extensive genomic data have been proposed [26,27], the overall distribution and frequency of nitrogen fixation in cyanobacteria remain poorly understood.

Identifying nitrogen-fixing potential from genomic data requires determining the presence of the six essential genes: *nifHDKENB* [28]. However, this is challenging for two reasons. First is the presence of nitrogenase-like enzymes (NFLs) [29]. Due to high amino acid sequence similarity, NFLs function other than nitrogen fixation, are often misidentified as *nif* genes [30]. In particular, cyanobacteria carries the dark-operative protochlorophyllide reductase (DPOR), an NFL involved in chlorophyll biosynthesis (Fujita and Bauer 2003)[31]. Thus, distinguishing *nifHDK* from the DPOR genes *chlLNB* is critical. Accurately discriminating between the similar gene pairs *nifDK* and *nifEN* is also essential [28]. Some strains carry vanadium (V)-dependent nitrogenase as well as molybdenum (Mo)-type nitrogenase [32,33]. The second reason is gene fragmentation. In many heterocystous cyanobacteria, *nif* genes are interrupted by long insertion sequences, complicating their identification. In *Anabaena* sp. PCC 7120, *nifD* is split by an 11-kb insertion that is precisely excised during heterocyst differentiation to reconstruct the functional *nifD* gene [34–37]. Such fragmentation is common among heterocystous species; for instance, in *Calothrix* NIES-4101, four genes (*nifBHDK*) are divided into 13 fragments, yet the functional genes are reconstructed in heterocysts, allowing for diazotrophic growth [38]. A previous genomic survey reporting only 87% of heterocystous cyanobacteria possessed the *nif* genes [12]might overlook some of such fragmented *nif* genes.

In this study, we determined the nitrogen-fixing potential of 586 cyanobacterial strains spanning diverse lineages with high-quality genomic data based on the conservation of *nifHDKENB* using a unique 2D-Similarity plot. Nearly all (97%) Nostocaceae strains were evaluated to be nitrogen-fixing indicating that this method is the most accurate to date. We developed Nif-Finder using these results as reference dataset, we evaluated the nitrogen-fixing potential in broader genomes from public genome collection including uncultured strains (2,718 strains), and found that Cyanobacteria phylum is one of the highest frequencies of nitrogen-fixing strains among 16 phyla. Furthermore, we identified that 21 genomes in several families carry “ancient” *nif* genes derived from anaerobic bacteria, rather than the “canonical” *nif* genes found in most cyanobacteria. We discussed the dynamic evolution of cyanobacterial *nif* genes in the context of their habitats, leading to a higher frequency of nitrogen-fixing strains than those in other bacterial phyla. (1,113 words)

## Results

### Identification of nif Genes

Prediction of nitrogen-fixing capability requires the accurate identification of the minimal set of nitrogenase genes: *nifHDKENB* [28]. In this study, we analyzed 586 high-quality cyanobacterial genomes obtained from NCBI and GTDB. Among them, 17 and 5 strains were classified into the non-photosynthetic sister groups of Cyanobacteria, Vampirovibrionia and Sericytochromatia, respectively. The remaining 564 strains belong to the Cyanobacteria, covering 18 diverse orders and 45 families (Supplementary Table S1). To distinguish NifHDKENB orthologs from Nif-like (NFL) sequences, cyanobacterial Nif HMM profiles that reflect the variation among cyanobacterial lineage were constructed. Using these HMMs as queries, Nif orthologs were searched and plotted on the 2D-Similarity plots (Fig. 1). Nif orthologs showed high sequence homology across the entire length of the HMM profile, resulting in the highest –log_10_(e-value) values and clustering near the full length of known Nif proteins on the x-axis (dashed circles in Fig. 1). These hits also matched the annotations in SWISS-PROT (Fig. 1), supporting accurate ortholog extraction. In contrast, NFL hits showed significantly lower –log_10_(e-value) values and were clearly separated from the Nif orthologs. To validate the prediction, maximum likelihood (ML) phylogenetic tree inferences were performed with all hit proteins. All assigned Nif orthologs formed monophyletic clades with known Nif sequences supported by high bootstrap values, while sequences identified as NFLs were placed in distinct clades (Supplementary Fig. S1). The results demonstrate that the 2D-Similarity plot estimation effectively distinguishes the cyanobacterial NifHDKENB orthologs from NFLs.

**Figure 1.**
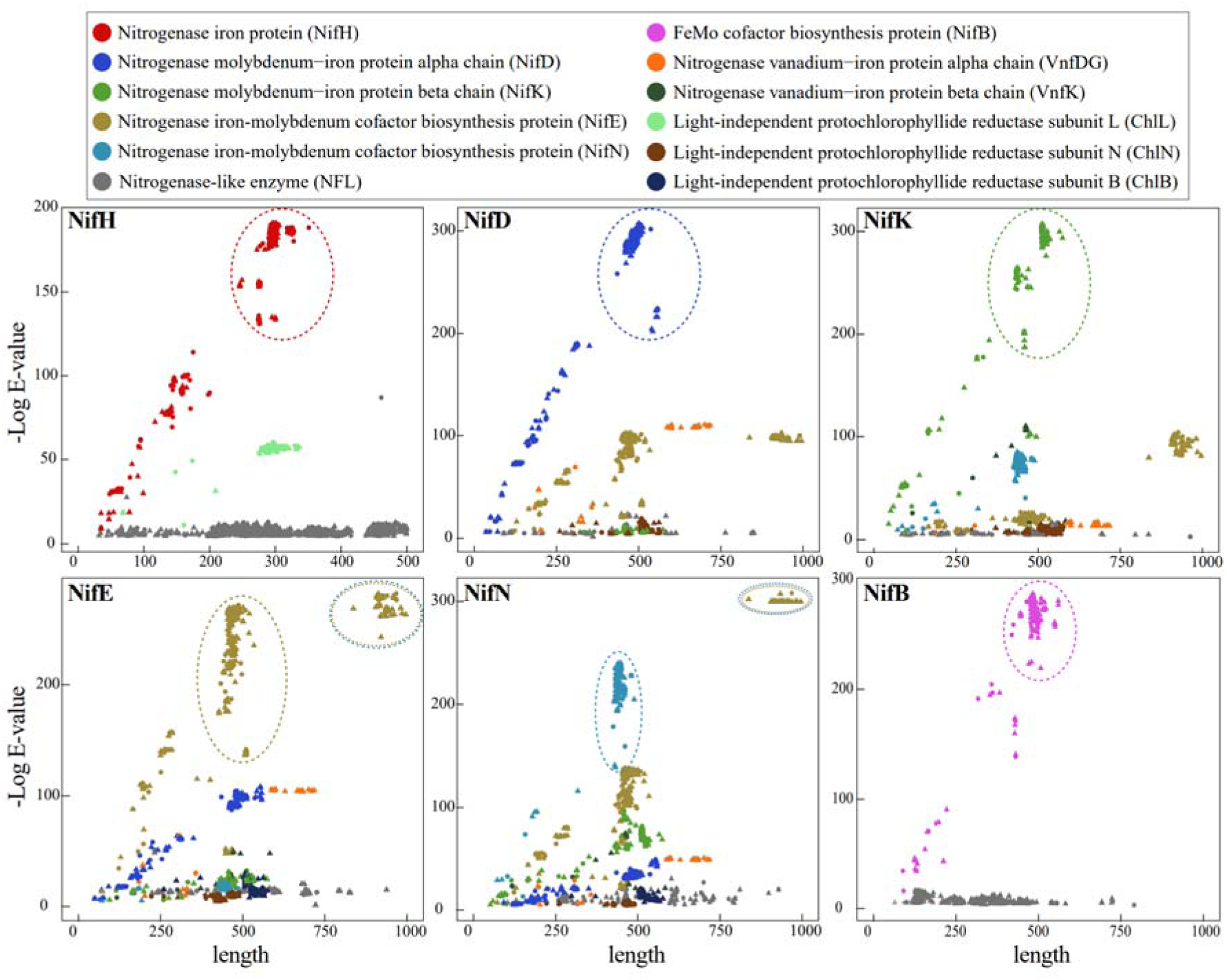
2D Similarity Plot of homology search for the six Nif proteins encoded by *nifHDKENB* of 586 cyanobacterial genomes. The relationship between –log_10_(E-value) of the Hmmer3 search using *nif* HMM profiles and the length of the hit protein against proteomes of 586 cyanobacterial strains are plotted. The color indicates the single best hit protein obtained by the BLASTp search against SWISS-PROT database. Circle plots represent hit from complete genome assembly, while triangle plots represent hits from draft genome assembly. The dashed circle indicates full-length hits of the target Nif. The double dashed circle represents the hits of the NifE-NifN fusion proteins.

In the 2D-similarity plots, many short hits appeared along a diagonal line from the origin to the Nif ortholog plots, indicating the presence of significantly shorter Nifs compared with full-length Nifs (Fig. 1). For example, in the NifH plot, unusual 142 short hits were identified, with lengths ranging from 10 amino acid residues (aa) to nearly the full length (270–300 aa). These short Nif hits exhibit four features as follows: 1) they are exclusively found in the family Nostocaceae, 2) they are detected in both draft and complete genomes, 3) they are located in close proximity to each other on the same contig, and 4) the short Nif sequences can be assembled to the full-length Nif sequence without overlaps or gaps. We concluded that these short hits are derived from fragmented *nif* genes by insertion sequences in the vegetative cell genomes [34]. The prevalence of short hits in all NifHDKENB proteins suggests that fragmentation by insertional sequences are common in the *nif* genes among Nostocaceae [37]. Indeed, we recently confirmed that in *Calothrix* sp. NIES-4101, despite the four *nif* genes, *nifH1DKB*, being fragmented into 13 parts, they are reconstructed precisely in heterocysts, enabling nitrogen-fixing growth [38]. Based on these findings, we determined the copy number of the nif genes in Nostocaceae, considering such fragmentation (e.g., if an open reading frame for *nifH* was found to be fragmented into three parts, it was estimated as a single copy of the *nifH* gene).

### Distribution of Nitrogen Fixation Capacity in Phylogenic Tree of Cyanobacteria

To elucidate the distribution of strains carrying nitrogen fixation capacity across cyanobacterial lineages, we mapped the presence of *nifHDKENB* on the phylogenetic tree constructed with 586 cyanobacterial strains, which includes 45 families and 219 genera level classification. The phylogenetic tree was constructed by the maximum likelihood (ML) method with 120 universal single-copy maker genes. It was rooted with Vampirovibrionia and Sericytochromatia using as outgroups (Fig. 2A). The presence or absence of *nifHDKENB* is indicated by red and black, respectively, in the outermost ring of the tree together with red and black dashed lines, respectively. The copy numbers of *nif* including *vnf* encoding V-nitrogenase, the presence of *nif*-related genes (*vupABC*, *patS*, *hetR*, *rbcL*, and *psbA*) and the presence of fragmented *nif* genes were plotted along the tree. No strains encoding *nifHDKENB* were found in the outgoups and the earliest-diverging genus *Gloeobacter* in the family Gloeobacteraceae (Fig. 2A). This is consistent with the previous study [39,40]. The *nifHDKENB* genes occur in all five strains in the genus *Thermostichus* of the family Thermostichaceae, which diverged after Gloeobacteraceae. This genus includes strains isolated from hot springs, such as *Synechococcus* JA-3-3Ab (Yellowstone A-Prime) and *Synechococcus* sp. JA-2-3B′a (2–13) (Yellowstone B-Prime). These strains are considered to represent lineages close to the origin of *nif* genes in cyanobacteria [41], which is consistent with our phylogenetic analysis.

**Figure 2.**
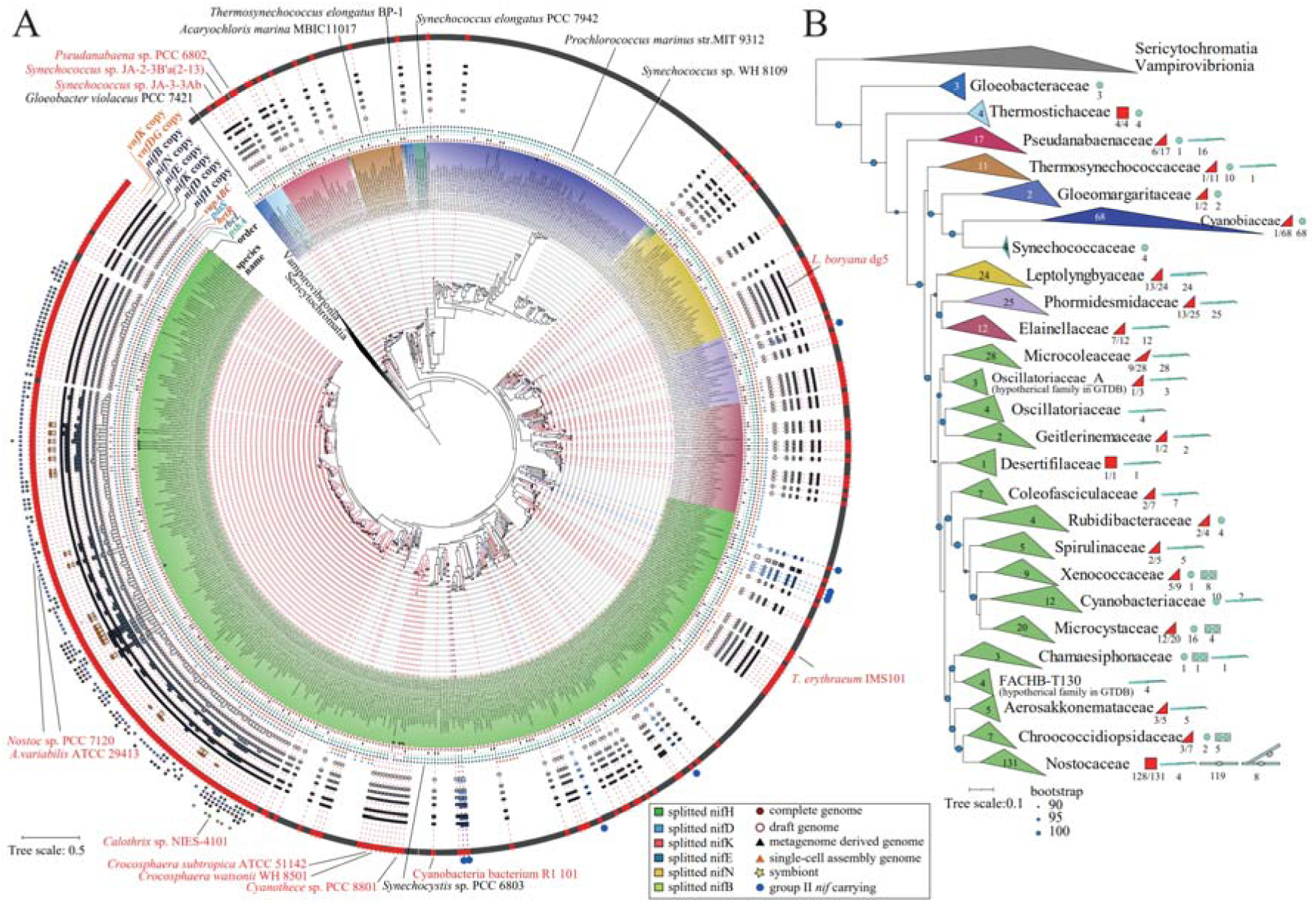
Maximum-likelihood (ML) phylogeny of cyanobacterial lineages showing the presence or absence of *nifHDKENB*. (**A**) Circular ML tree of 586 cyanobacterial lineages. The tree was inferred using the ML method based on 120 genes conserved across bacteria. The presence or absence of *nifHDKENB* is indicated by red and black, respectively, in the outermost ring together with red and black dashed lines, respectively, connecting the tree and the species names. The blue small circles just outside of the outermost ring indicate genomes encoding ancient *nif* genes (Group II). Copy numbers of *nifHDKENB* and *vnfDKG* are shown in the outer small box plots (gray for MoFe-type *nifHDKENB*; orange for VFe-type *vnfDKG*). The fused *nifEN* was determined to be redundant, with one copy in each of *nifE* and *nifN*. Additionally, information on the fragmented *nif* genes is shown in small boxes just outside the outermost ring, and the presence or absence of related genes (*vupABC*, *patS*, *hetR*, *rbcL*, and *psbA*) is shown in small circles inside the ring denoting the copy numbers of *nifHDKENB*. Representative species names (strains with *nifHDKENB* in red and those without in black) are shown just outside the outermost circle for easy identification. (**B**) Rectangular ML tree collapsed at the family level. Twenty-six major families of Oxyphotobacteria from the circular tree are shown. The number of strains with known morphologies is plotted along with small icons showing morphology at each clade. Blue circles on the branches represent bootstrap values of 95-100 based on 1,000 ultrafast bootstrap replicates implemented in IQ-TREE 2. Sericytochromatia and Vampirovibrionia were used as outgroups. The full-size figure is available from the Nif-Finder repository.

Regarding lineages diverging after *Thermostichus*, we first focused on heterocystous strains. These strains belonging to the family Nostocaceae formed a monophyletic group, which is consistent with previous studies. In this clade, 97.0% of strains (163 out of 168) were predicted to have the nitrogen fixation capacity (Fig.2A). With the exception of five strains (Supplementary Table S5), the result confirmed the close association between heterocyst formation and nitrogen fixation capacity. Furthermore, our analysis revealed that *nif* fragmentation is widespread across most genera in the family Nostocaceae, including *Nostoc*, *Dolichospermum*, *Nodularia*, *Trichormus*, *Aulosira*, and *Calothrix* (Fig. 2A). Fragmentation was detected in at least one of the *nifHDKENB* genes. Among them, *nifD* exhibited the highest frequency of fragmentation (122 of 163 strains), followed by *nifH* (64 strains), *nifK* (37 strains), *nifE* (21 strains), *nifB* (7 strains), and *nifN* (1 strain). In contrast, the *Raphidiopsis* strains showed no *nif* fragmentation (see Discussion).

Among non-heterocyst-forming cyanobacteria (182 strains), the distribution of *nifHDKENB* exhibited a highly mosaic pattern, indicating a scattered distribution of nitrogen fixation capacity (Fig.2A). To examine the relationship between nitrogen fixation and morphology, morphological information was curated from the literature for all strains. The 182 strains were classified into unicellular (Section I, 84 strains), colonial/beaocyte-forming (Section II, 8 strains), and filamentous (Section III, 90 strains) groups (Table 1; Supplementary Table S1). As in previous studies [42], filamentous morphology was polyphyletic. The frequency of nitrogen fixation (FNF) was estimated to be 22.6% in unicellular strains (19 of 84), 62.5% in colonial strains (5 of 8), and 44.4% in filamentous strains (40 of 90). Compared with multicellular strains, unicellular strains showed a lower FNF, suggesting the difficulty of simultaneously performing oxygenic photosynthesis and nitrogen fixation in unicellular forms.

**Table 1.** Distribution of *nif* genes across the five cyanobacterial Sections I–V.

| Section <sup>*1</sup> | Morphology <sup>*1</sup> | Number of genomes |  | Frequency of nitrogen fixation (FNF, %) |  |
| --- | --- | --- | --- | --- | --- |
|  |  | subtotal <sup>*2</sup> | Potential Diazotroph |  |  |
| I | Unicellular | 84 | 19 | 22.6 | 35.2 |
| II | Unicellular or baeocytes | 8 | 5 | 62.5 |  |
| III | Filamentous | 90 | 40 | 44.4 |  |
| IV | Filamentous | 69 | 67 | 97.1 | 97.4 |
| V | Filamentous with true/pseudo branching | 9 | 9 | 100.0 |  |
| Total |  | 260 | 140 | 53.8 |  |

<sup>\*1</sup> The classification was based on the definitions in Rippka et al. 1979.
<sup>\*2</sup> Calculations were only performed for strains (260 strains) whose morphology was determined from literatures.

Unicellular nitrogen-fixing lineages were primarily classified into the families Thermostichaceae (including *Synechococcus* JA-3-3Ab) and Microcystaceae (including *Synechocystis* sp. PCC 6803 and *Crocosphaera subtropica* ATCC 51142). All five Thermostichaceae strains and 12 of 20 Microcystaceae strains carried *nifHDKENB* (Fig. 2B). In contrast, most strains within the family Cyanobiaceae, which includes the marine genera *Prochlorococcus* and *Synechococcus*, lacked *nifHDKENB* (67 of 68 strains), consistent with the extensive genome reduction observed in this family [43]. The only exception is *Vulcanococcus limneticus* LL. However, diazotrophic growth was not observed in this strain [44].

Compared with unicellular strains (Section I), strains in Sections II and III showed higher FNF values, indicating a more mosaic distribution of the *nifHDKENB* genes. Interestingly, mosaic patterns were observed even in well-characterized diazotrophic genera, such as *Chroococcidiopsis* (Section II), *Symploca*, *Oscillatoria*, *Pseudanabaena*, and *Leptolyngbya* (Section III) [24,45–49]. This observation suggests that independent losses of the *nif* genes have frequently occurred across cyanobacterial lineages during speciation (Supplementary Table S3).

Overall, the *nifHDKENB* genes were detected in 285 out of 586 strains, resulting in an FNF of 48%. This value is consistent with the current consensus that about half of cyanobacterial species possess diazotrophic capacity.

### Distribution of nif Genes in Cyanobacteria and Other Phyla

To gain a more comprehensive understanding of distribution of nitrogen fixation capacity in cyanobacteria, we developed a python script “Nif-Finder” for identifying *nifHDKENB* genes to enable large-scale analyses. This script automatically detects the NifHDKENB proteins while discriminating from NFLs. Using the results obtained from the 586 strains as a reference dataset, this script determines the annotation of a sequence based on the nearest distance within the 2D-Similarity plot. Nif-Finder was applied to cyanobacterial 1,365 species-level clades (ANI < 95%) in NCBI GenBank assembly [50], revealing that 498 clades (FNF, 36.4%) carrying at least one copy of *nifHDKENB* were predicted to have nitrogen fixation capacity (Table 2). Nif-Finder was also applied to GTDB representative genome (R226; 2,718 genomes), which incorporates a larger proportion of single-cell and metagenome-assembled genomes (MAGs) [51]. Only 527 (FNF, 19.4%) were predicted to have nitrogen fixation capacity (Table 2). A clear trend was observed, in which the proportion of strains with nitrogen fixation capacity significantly decreased as the ratio of uncultivated strains in the analyzed strains increased. This trend implies that the proportion of cyanobacterial strains capable of nitrogen fixation in field environments is significantly lower than expected based on the current knowledge of isolated and cultured cyanobacterial strains.

**Table 2.**
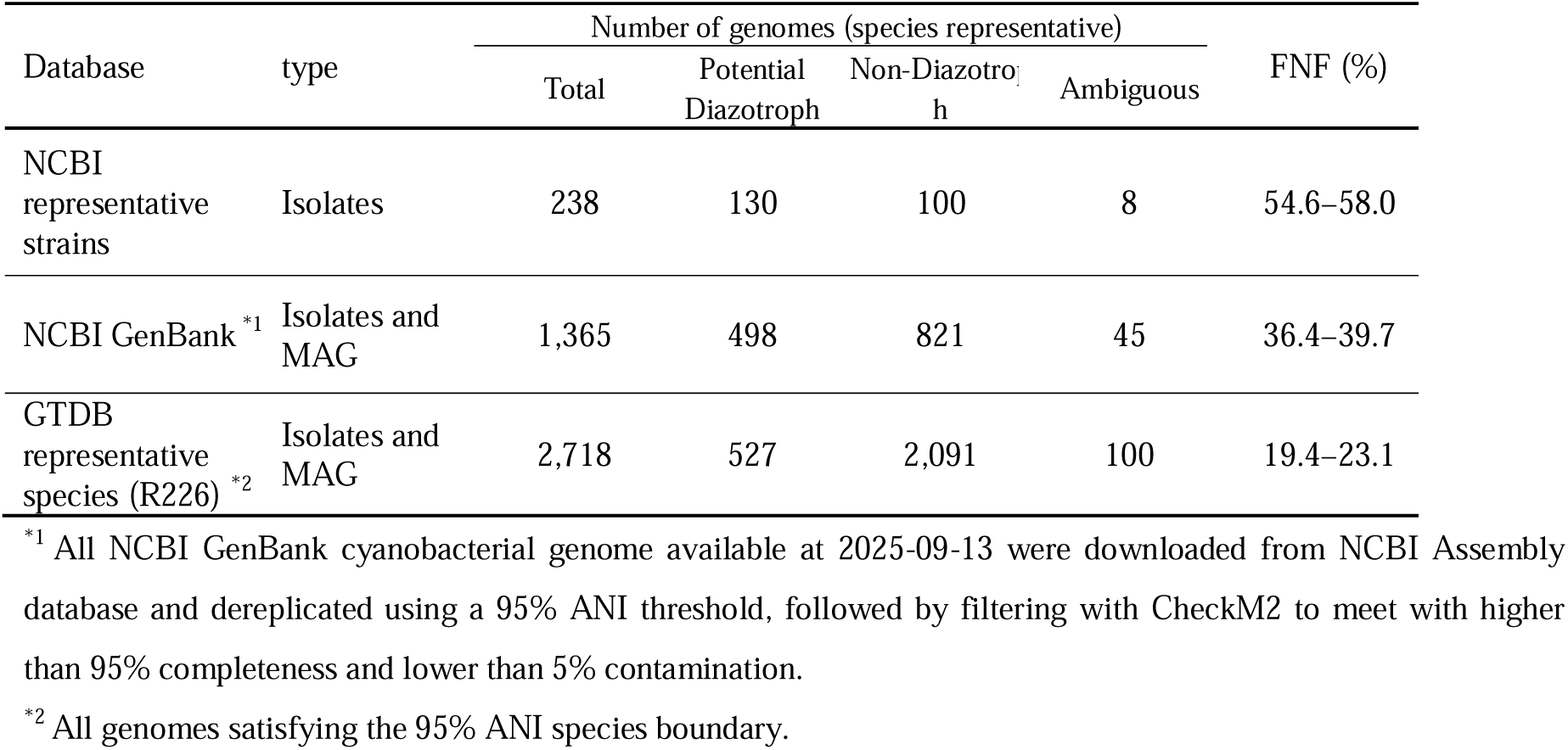
Prediction of nitrogen fixing capacity of cyanobacteria using Nif-Finder.

We compared FNF values in cyanobacteria with those of other phyla in the GTDB resource (Fig. 3). Excluding Ambiguous strains, the Cyanobacteria (19.4%) showed the highest FNF value, followed by Nitrospirota (11.8%), Campylobacterota (10.4%), Pseudomonadota (Proteobacteria) (5.7%), and Thermodesulfobacteriota (5.1%). Other major phyla, such as Bacillota (Firmicutes) and Actinomycetota (Actinobacteria), exhibited much lower FNF values of 0.4% and 0.1%, respectively. Including ambiguous strains, the Cyanobacteria phylum ranks fourth or third (excluding phyla with fewer than 100 strains). However, these results demonstrate that Cyanobacteria is the phylum exhibiting one of the highest FNF values among all bacterial phyla.

**Figure 3.**
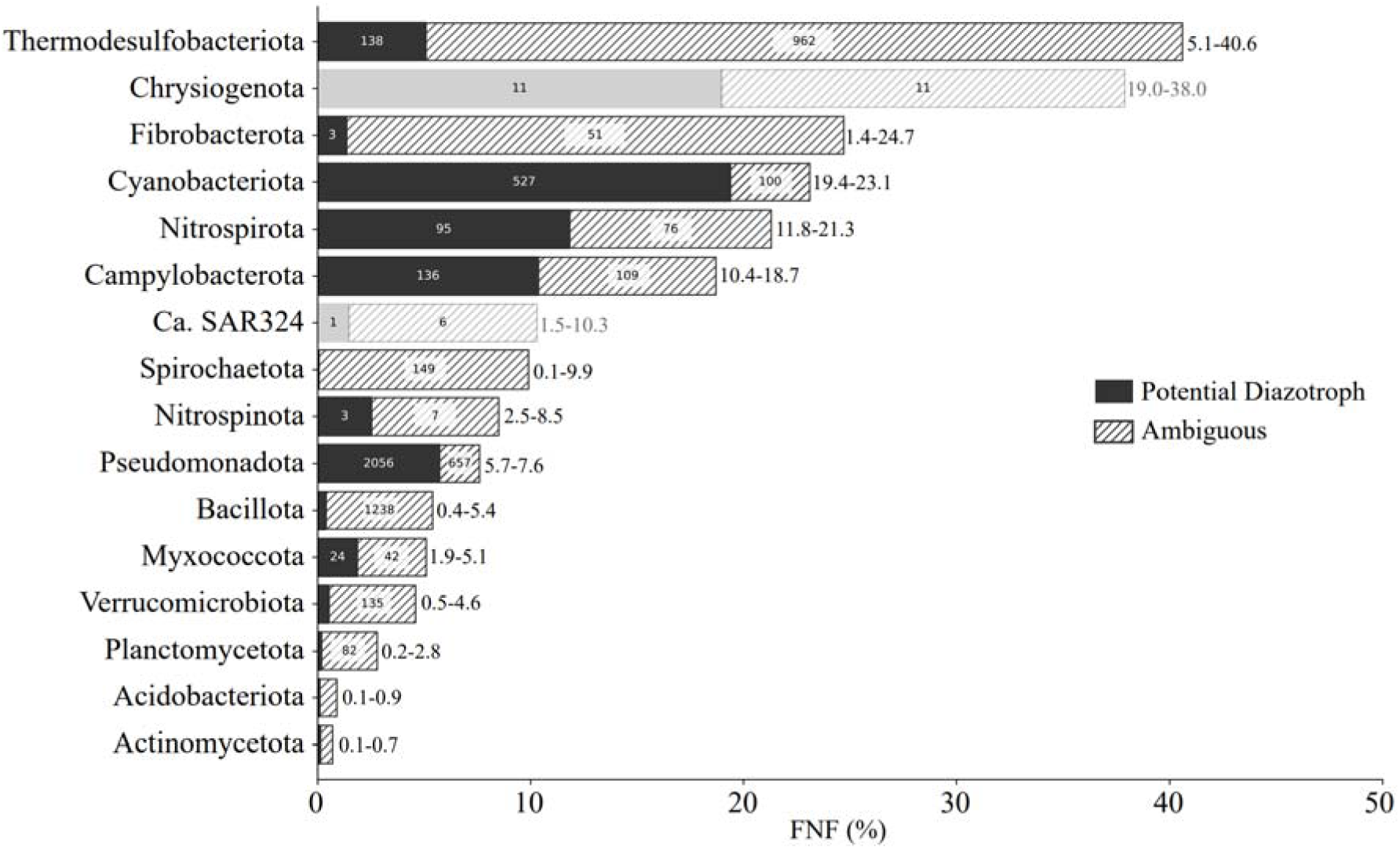
The frequency of nitrogen-fixing (FNF) strains (*nifHDKENB* carrying strains) across phyla. The FNF values (%) were estimated in the 110 phyla based on 143,619 representative bacterial genomes from GTDB R226 and top 16 phyla are shown. The bar represents the maximum value of FNF (%) consisting of solid and hatched bars, which indicate FNF strains carrying the full set of *nifHDKENB* (Potential Diazotrophs) and those carrying only a partial set (Ambiguous), respectively. Actual % values of Potential Diazotrophs and the sum of Potential Diazotrophs and Ambiguous strains are shown in just right of the bars. Numbers of the genomes are shown inside the bars. Phyla with a total genome number (n) of 100 or less are shown in lighter colors considering insufficient sample number.

### Identification of Group II nif

During the screening of 1,365 cyanobacterial genomes using Nif-Finder, 45 genomes (3.3%) were identified as lacking one or more *nifHDKENB* genes. Interestingly, 21 of 45 commonly lacked only *nifN*. These strains were restricted to four specific genera from different families: *Coleofasciculus*, *Roseofilum*, *Sodalinema*, and *Hydrococcus*. Due to an unexpectedly low amino acid identity (approximately 35%) to the known NifN sequences, their NifNs of the 21 strains were initially misidentified as NFL or NifK (Supplementary Figure S3).

NifDK and NifEN are thought to have evolved via gene duplication from a common ancestor followed by divergence into NifD/NifE and NifK/NifN pairs [52–54]. Molecular phylogenetic analysis showed that these NifN-like sequences branched shortly after the divergence of NifK and NifN (Supplementary Fig. S5). Similarly, the NifE sequences in these genera also diverged shortly after the divergence of NifD and NifE. A similar trend was observed in NifH and NifB tree (Supplementary Fig. 6). Further screening using early-diverging NifEN-like sequences as queries revealed that these types of Nif genes are present in a total of 10 cyanobacterial genera, including *Baaleninema* and five GTDB-defined hypothetical genus level classification (CCY1219, JANQEB01, LEGE-06147, JAQMSD01, and PCC-9228).

Raymond et al. proposed a classification (Groups I to V) for nitrogenases (Mo-, V-, and Fe-type) and NFLs [55], in which Mo-nitrogenases form Groups I and II. Group I consist of “canonical Nif” found in aerobic and facultative anaerobic bacteria (e.g., *Azotobacter vinelandii*, *Klebsiella pneumoniae*, rhizobia, and purple bacteria) including most cyanobacterial *nif* genes [4]. Group II includes nitrogenases of obligate anaerobes such as *Clostridium acetobutylicum*, *Chlorobium tepidum* and methanogens, which diverges closer to the root (Fig. 4). We refer the Group II Nif to “ancient Nif”. Multiple sequence alignment of Nif proteins showed some unique features of ancient Nif while the conserved cysteine residues to chelate the metallocenters (Supplementary Figs. 4 and 8) [56,57]. Our re-evaluation of the genera harboring these unique NifEN sequences showed that they belong to Group II and are distinct from the Group I Nif found in most cyanobacteria. This finding demonstrates that some cyanobacteria genera possess ancient Nif (Group II) rather than the canonical Nif (Group I) [12,58].

**Figure 4.**
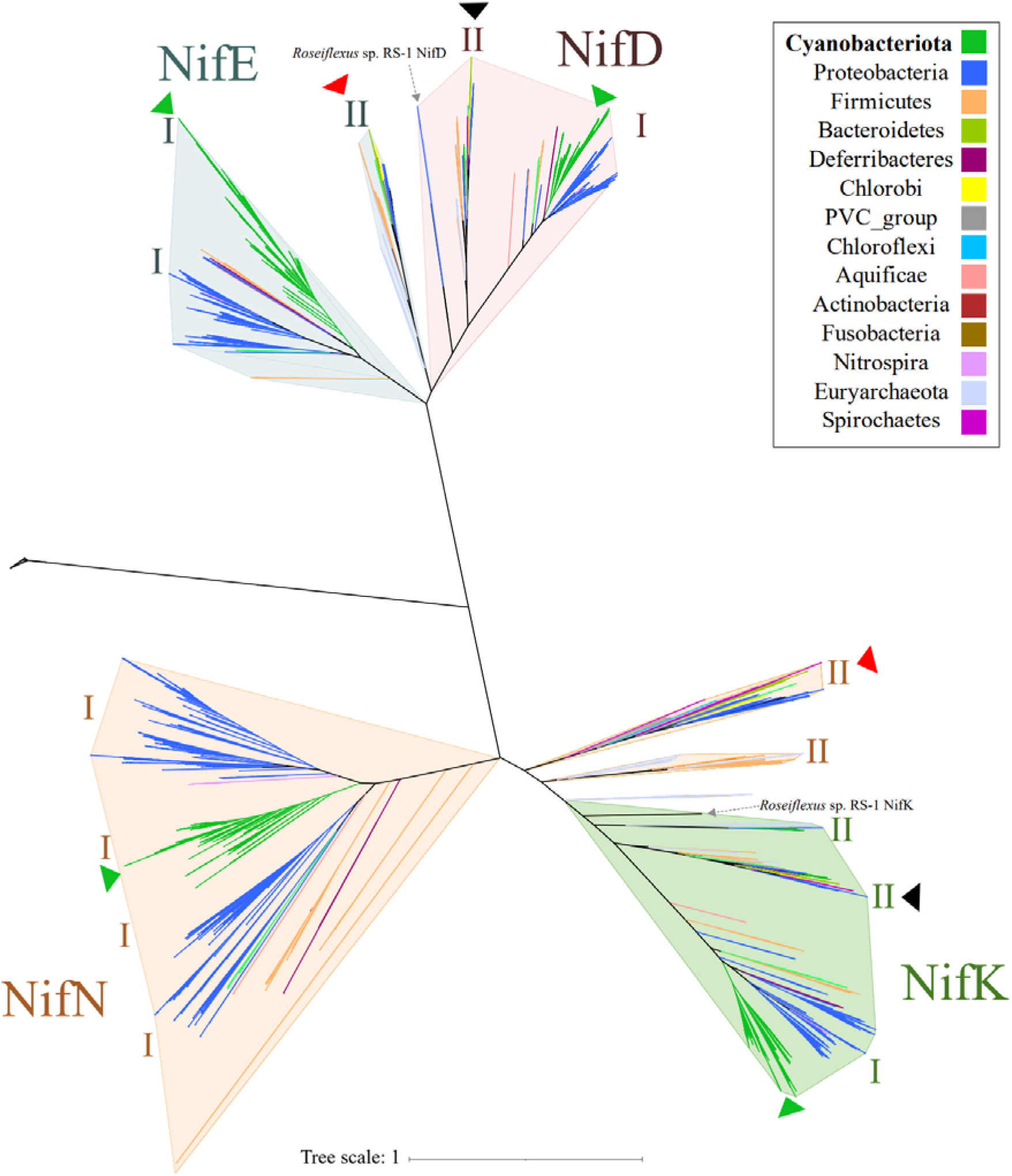
Unrooted phylogenetic tree of the four proteins, NifD (light pink), NifK (pale green), NifE (light blue) and NifN (light orange). NifDKEN sequences from 1,365 cyanobacterial species-level representative genomes and 853 genomes from strains belonging to other bacterial phyla (<u>Koirala and Brözel</u>, 2021) were used. Green triangles indicate the cyanobacterial clades with the canonical Nifs belonging to Group I. Red triangles indicate the clades including the cyanobacterial ancient NifE and NifN (Group II) together with anaerobic bacteria and a subset of methanogenic archaea. Black triangles indicate the NifD and NifK clades belonging to Group II, which also diverged early in the NifD and NifK proteins. NifDK from *Roseiflexus* sp. RS-1, which was shown to be functional nitrogenase without NifEN (<u>Payá-Tormo et al. 2024</u>), was also shown by dotted arrows as a representative of Group II.

### Comparison of nif Gene Clusters

In most diazotrophs including nitrogen-fixing cyanobacteria, the *nifHDKENB* genes, are included in a compact *nif* gene cluster together with other accessory and *nif*-related genes [28],[59]. Cyanobacterial *nif* cluster frequently includes the *cnfR* gene encoding the master regulator of the *nif* genes. CnfR activates the transcription of the *nif* genes by recognizing *cis* elements upstream of *nifB* and *nifP* [48]. Given the presence of the ancient Nif in some cyanobacteria, we compared the gene arrangement of the *nif* gene clusters encoding the canonical and ancient Nifs. In strains harboring the canonical Nifs, the core *nif* gene cluster consists of the nine genes including *nifHDKENB*: *nifB-nifS-nifU-nifH-nifD-nifK-nifE-nifN-nifX* (Fig. 5). While the gene order is highly conserved, some variations occur; for instance, in *Leptolyngbya boryana* dg5, *nifE-nifN-nifX* is encoded as a sub-cluster on the opposite strand. Accessory *nif* genes such as *nifP*, *nifV*, *nifW*, *nifT*, and *nifZ*; and *nif*-related genes such as *hesAB–fdxB–feoA*, *desA*, *mop*, and *modABC*; as well as unknown conserved ORF such as *orf99* and *orf155* (Tsujimoto et al. 2014)[48] are located flanking the core *nif* genes (Fig. 5). The average number of genes within the canonical *nif* cluster is 25, which is significantly higher than the 15 in the model diazotroph *A. vinelandii* [60]. This suggests that cyanobacteria, which solve the challenge of protecting nitrogenase from oxygenic photosynthesis and aerobic environment, require the additional genes and the strictly coordinated regulation of *nif* expression.

**Figure 5.**
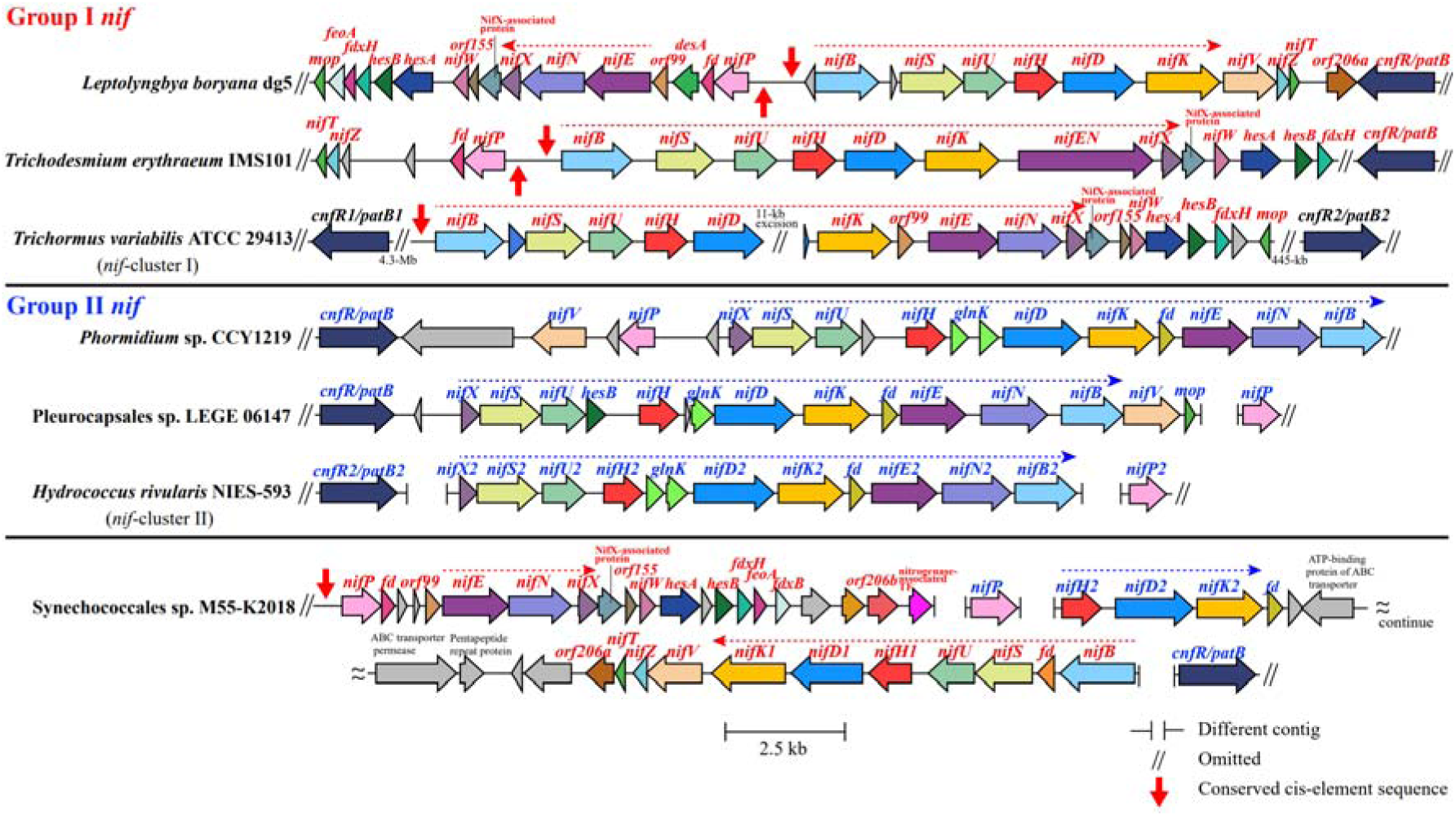
Comparison of the *nif* gene clusters of Groups I and II *nif* genes in cyanobacteria. ORFs are shown as thick horizontal arrows, with identical colors indicating orthologous genes. Gene names for Groups I and II are highlighted in red and blue, respectively. Red and blue horizontal dotted lines indicate the conserved core *nif* cluster genes in Groups I and II, respectively. *Synechococcales* sp. M55-K2018 encodes both Group I (canonical) *nifHDKENB* and Group II (ancient) *nifHDK*. Red vertical arrow indicates the conserved *cis*-element sequences recognized by the CnfR transcriptional regulators directly involved in the transcriptional activation of the *nif* gene cluster (<u>Ryoma Tsujimoto et al., 2016</u>).

The ancient *nif* gene clusters exhibited a distinct gene composition and an order compared with that of the canonical *nif* gene clusters. The typical core consists of 10 genes: *nifS-nifU-nifH-glnK-glnK-nifD-nifK-nifE-nifN-nifB*. The *nifB* is located at the last position, and two *glnK* genes (encoding P_II_ proteins) located in tandem between *nifH* and *nifD* (Fig. 5). The average number of genes in the ancient *nif* clusters was only 15 (Fig. 5). Several genes that are highly conserved in the canonical nif clusters, including *orf99*, *nifW*, *hesAB*, *nifZT*, and *cox*, were absent from the ancient nif clusters (Table 3). The *cnfR* gene is found within or in the neighborhood of the ancient *nif* cluster, however, the *cis*-elements conserved upstream of *nifB* in the strains carrying the canonical *nif* genes [48] were not found. This feature suggests that the regulatory mechanism of the ancient *nif* gene clusters differ from that of the canonical *nif* gene cluster. The order and orientation of the cyanobacterial ancient *nif* cluster resemble those of the anaerobic bacteria. *Desulfomonile tiedjei* DSM 6799 and *Chlorobaculum tepidum* TLS, which represent the closest strains in the NifH phylogenetic tree to the cyanobacterial ancient NifH (Supplementary Fig. 7), shared an identical arrangement of the core genes; *nifH–glnK–glnK–nifD–nifK–nifE–nifN–nifB*, just missing *nifUS* (Supplementary Fig. S8).

**Table 3.** Conservation of the genes in the canonical (Group I) and ancient (Group II) *nif* clusters across potential nitrogen fixing cyanobacterial strains.

| Gene<sup>*1</sup> | Presence in canonical<br>*nif*-cluster from 65<br>strains | Presence in ancient<br>*nif*-cluster from 11<br>strains | Gene<sup>*1</sup> | Presence in canonical<br>*nif*-cluster from 65<br>strains | Presence in ancient<br>*nif*-cluster from 11<br>strains |
| --- | --- | --- | --- | --- | --- |
| *nifH* | 65 | 11 | *nifP* | 46 | 3 |
| *nifD* | 65 | 11 | *orf99* | 42 | 0 |
| *nifK* | 65 | 11 | *fdx* | 41 | 11 |
| *nifE*<sup>*2</sup> | 65 | 8 | *fdxB* | 41 | 0 |
| *nifN*<sup>*2</sup> | 65 | 8 | *feoA* | 39 | 0 |
| *nifB* | 65 | 9 | 4Fe-4S ferredoxin | 36 | 0 |
| *nifU* | 65 | 9 | *cnfR* | 30 | 5 |
| *nifS* | 65 | 9 | *modA* | 26 | 3 |
| *nifX* | 65 | 9 | *orf206a* | 26 | 0 |
| *nifW* | 65 | 0 | *modBC* | 25 | 3 |
| *deoA* | 64 | 0 | *desA* | 24 | 0 |
| *orf155* | 63 | 0 | *tetR* | 24 | 0 |
| *orf70* | 63 | 0 | *tam* | 24 | 0 |
| *hesA* | 63 | 0 | CBS<br>domain-containing<br>protein | 22 | 0 |
| *hesB* | 59 | 2 | *arsC* | 19 | 0 |
| *mop* | 58 | 6 | *fdxH* | 18 | 0 |
| *nifZ* | 57 | 0 | *feoB* | 16 | 0 |
| *nifT* | 57 | 0 | *fldA* | 10 | 0 |
| *nifV* | 46 | 6 |  |  |  |

<sup>\*1</sup> Gene names *orfxxx* are from *L. boryana* dg5 (Tsujimoto et al. 2014).
<sup>\*2</sup> The *nifE-nifN* fusion gene was counted redundantly.

### Cyanobacteria Harboring Both Canonical and Ancient nif Clusters

We found cyanobacterial strains that carry both canonical and ancient *nif* clusters (Fig. 5). These strains were classified into three types. First, three strains in *Hydrococcus* carry the complete sets of both ancient and canonical *nif* gene clusters. Second, one strain of *Limnospira* (*L.* sp. CCY1209) exhibited a hybrid feature, containing ancient *nifHDKB* and canonical *nifEN*, which constitute the full minimal *nif* gene set. Third, 38 strains of *Microseira* carried a canonical *nifHDKENB* and a duplicated ancient *nifHDK* (e.g., *M. wollei* NIES-4236, *M. sp.* BLCC-F43, Supplementary Fig. S8). A similar arrangement was observed in a Synechococcales cyanobacterium M55-K2018 (GTDB genus *DVEB01*), where the ancient *nifHDK* was encoded adjacent to the canonical *nif* cluster (*nifBSUHDK*) (Fig. 5). In total, 48 strains harboring both canonical and ancient *nif* clusters were distributed across four genera (Supplementary Table S11). Cyanobacteria carrying the ancient *nif* clusters are thought to have acquired these genes via horizontal gene transfer (HGT) from some anaerobic bacteria. These cyanobacteria may functionally utilize the ancient *nif* for nitrogen fixation under strictly anaerobic conditions, such as microbial mats, while employing the canonical *nif* in oxic environments. Such functional differentiation of the two *nif* types would provide a selective advantage. Or the coexistence may represent an ongoing evolutionary transition toward the convergence of a single *nif* system by losing either one.

### Environmental Distribution of Cyanobacteria Harboring Ancient nif Clusters

Cyanobacteria carrying the ancient *nif* genes may specifically inhabit some anaerobic environments. To examine this possibility, publicly available metagenomes and their correlations with environmental niches were analyzed. Twenty major cyanobacterial niches were defined (freshwater, water reservoirs, eutrophic water, freshwater sediment, soda lake, hot spring, brackish water, salt marsh, marine, coral reef, soil, field, rhizosphere, phyllosphere, moss, antarctica, glacier, biofilm/mat, phycosphere, and wetland/paddy-field). From over 1.26 million publicly available metagenomes, datasets corresponding to each keyword were downloaded, and genus-level cyanobacterial abundance was quantified using exact k-mer matching with Kraken2. Hierarchically clustered heatmaps of average abundance across environments are shown in Figure 6, with the non-cyanobacterial niche “Fermentation” included as a negative control. Marine nitrogen-fixing cyanobacteria, such as *Trichodesmium* and *Crocosphaera* [61,62], were dominant in Marine and Coral-reef environments. *Nostoc*, which frequently associates with plants or moss [63], was mainly detected in Moss. These results are consistent with previous reports, supporting that our analysis reflects the global ecological niches of cyanobacteria. Among the seven genera carrying the ancient *nif*, the four genera *Roseofilum*, *Baaleninema*, *Microseira*, and *Coleofasciculus* were predominantly detected in Biofilm/Mat (Fig. 6). This niche overlaps with that of anaerobic bacteria, suggesting that physical proximity with anaerobic bacteria in such environments may have facilitated HGT of ancient *nif* genes [4].

**Figure 6.**
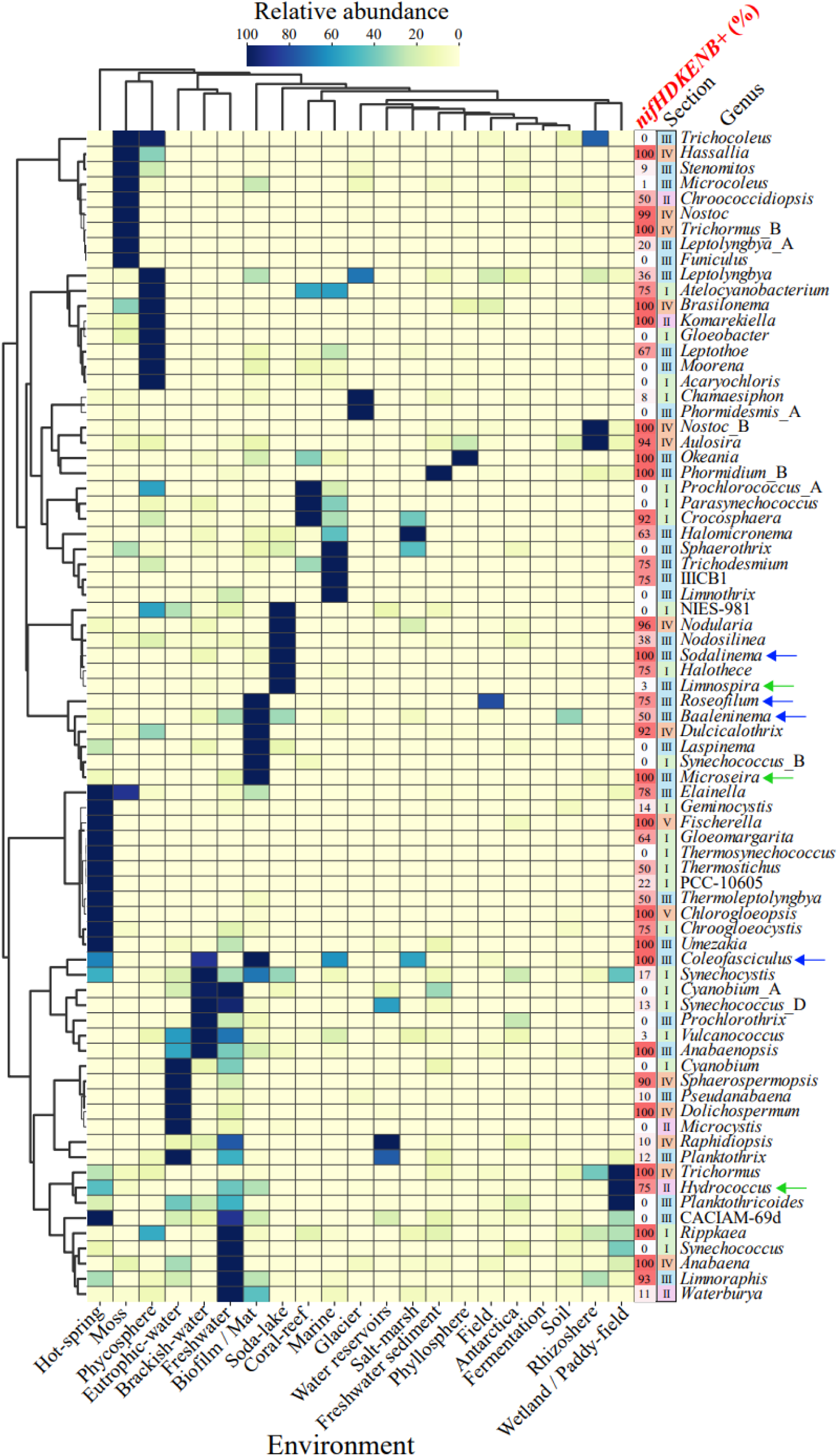
Metagenomic profiling of cyanobacteria at the genus level across diverse environments. Abundance profiles of 77 major cyanobacterial genera were investigated based on counts of 1.26 million publicly available metagenomic sequencing from the NCBI Sequence Read Archives. For each SRA/ERA/DRA dataset, 300,000 reads were subsampled and analyzed using the Kraken2 taxonomic profiler against a custom database of 2,716 GTDB R226 cyanobacterial representative genomes. Relative abundance below 0.1% were excluded, and average abundances were row normalized within each genus (scaled 0–100) to highlight distribution patterns across diverse environmental niches. Hierarchical clustering was performed using Ward’s method based on Euclidean distances. Sections I–V and FNF (%) per genus (*nifHDKENB*+) are displayed alongside the genus names. Four genera containing strains with the ancient *nif* (Group II) genes are shown with blue arrows and three genera containing strains with both canonical (Group I) and ancient (Group II) *nif* genes are shown with green arrows.

The genus *Hydrococcus*, which harbor both canonical and ancient *nif* clusters, were abundant in Wetland/Paddy-field environments alongside canonical *nif*-carrying genera, and were also detected in Hot-spring, Freshwater, and Biofilm/Mat environments, implying a correlation between the coexistence of both *nif* types and environmental conditions (Fig. 6). These findings suggest that under strictly anaerobic environments the ancient *nif* genes can be utilized for nitrogen fixation even performing oxygenic photosynthesis.

## Discussion

### Distribution of Nitrogen Fixation Capacity and the Development of Nif-Finder

In this study, we investigated the distribution of nitrogen fixation capacity across all cyanobacterial lineages by accurately predicting the presence or absence of the six marker genes: *nifHDKENB*. While nearly all heterocyst-forming cyanobacteria were identified as diazotrophs, nitrogen fixation capacity in non-heterocystous lineages was distributed in a mosaic manner. Based on these findings, we developed automated script: Nif-Finder, and applied it to a broader range of cyanobacteria, including uncultivated species and MAGs, to calculate the FNF. The resulting FNF of 19.4% demonstrates that the phylum Cyanobacteria possesses one of the highest proportions of nitrogen-fixing species (Fig. 3, Table 2). Our analysis also revealed that strains carrying the ancient (Group II) *nif* genes were distributed across multiple genera (Fig. 4). Additionally, we identified several strains that harbor both canonical and ancient *nif* systems simultaneously (Fig. 5).

### 2D-Similarity Plot: Accurate Way for Nif Identification

Accurate identification of the *nifHDKENB* genes from genomic data are challenging by two primary reasons: the discrimination of *nif* genes from NFL homologs (including the differentiation between NifDK vs. NifEN and Mo- vs. V-nitrogenases) and the presence of fragmented *nif* genes. We addressed these issues using a unique 2D-Similarity plot visualization (Fig. 1). In this plot, the vertical axis represents the -log of the E-value obtained from homology searches using custom profile HMMs, while the horizontal axis represents the length hit protein. Nif orthologs clustered in the upper-right region of the plot (indicated by dashed circles in Fig. 1), allowing for clear discrimination from NFLs, NifEN, or Vnf. An exception is the ancient-type NifN: because it branches near the NifN–NifK divergence node (Fig. 4), Group II NifN represents a case in which a clear distinction is difficult in our method.

### Detection of Fragmented nif Genes

The 2D-Similarity plot facilitates the identification of fragmented *nif* genes, as they appear as plots showing a linear distribution between the origin and the primary ortholog cluster. The *nif* fragmentation was first discovered in the heterocyst-forming cyanobacterium *Anabaena* sp. PCC 7120, where the *nifD* coding region is interrupted by an 11-kb insertion element [34–36]. The similar interruptions were subsequently found in the *fdxN* gene coding ferredoxin and the *hupL* gene coding hydrogenase. Systematic surveys have revealed that *nif* fragmentation occurs at a high frequency in heterocyst-forming cyanobacteria (Hilton et al., 2016)[37]. We identified one species *Calothrix* sp. NIES-4101 carrying extensively fragmented *nif* genes. In the genome, the *nifBHDK* are fragemented into 13 parts spanning a 350-kb region with different order and orientations. Nevertheless, a functional *nifBHDK* operon is restored via multi-step programmed genome rearrangements [38].

By accurately determining fragmented *nif* genes, we estimated the FNF of heterocyst-forming species to be 97.0%, which is 10% higher than the 86.7% reported by Chen et al. (2022)[12]. We hypothesize that automated approaches analyzing large datasets (e.g., 650 strains), including draft genomes, often fail to detect fragmented *nif* genes, primarily because *ab initio* gene prediction fails to identify such fragmented structures. To detect fragmented *nif* genes, Prodigal was run in both training and anonymous modes. In addition, six-frame BLASTx search was performed to detect very short *nif* fragments (encoding <20 amino acids). Fragmentation also occurs in *fdxN*, *hupL*, and *cox*; therefore, it is important to consider that other heterocyst-specific genes may likewise be fragmented in the genome assembly from vegetative cell (Hilton et al., 2016; Uesaka & Banba et al., 2024)[37,38].

### nif Copy Number Variation in Heterocyst-Forming Species

While the copy number of *nifHDKENB* was one in most strains, within the family Nostocaceae, strains with a second copy of *nif* were often found (Fig. 2A, Supplementary Fig. S10). Copy numbers were highest in *Aulosira* (average 2.3) and *Nostoc* (average 1.7) in the Nostocaceae genera. The *nifH* showed particularly high values, averaging 3.8 in *Aulosira* and 3.2 in *Nostoc*. These lineages frequently encode V-nitrogenase (*vnfDKG*), which is found in 6 of 10 *Aulosira* strains and 6 of 17 *Nostoc* strains and all members exhibited *nif* fragmentation (Fig. 2A; Supplementary Table S1). In contrast, *Raphidiopsis* strains carried a single *nif* copy and lacked both *vnfDKG* and fragmentated *nif* (Fig. 2A). A similar trend was observed in *Dolichospermum*, which is phylogenetically close to *Raphidiopsis* (Supplementary Table S1). Heterocyst-forming cyanobacteria tend to possess relatively large genomes, with 9.45 Mb of *Aulosira* and 8.67 Mb of *Nostoc* which are in the largest sizes in cyanobacteria [64]. In contrast, the two genera, *Dolichospermum* (5.3 Mb) and *Raphidiopsis* (3.7 Mb), have the smallest genomes within the Nostocaceae (Supplementary Figure S10). Consistent with this, three *Raphidiopsis* strains (D9, CENA303, and NIES-932) lacking the *nif* genes [65–67] have the smallest genome size, ranging from 3.19 to 3.43 Mb. The three strains may represent highly streamlined genomes [68]. Their features could be considered as an evolutionary model of stepwise genome streamlining. Such evolution involves loss of V-nitrogenases, decrease of the copy number of *nif* genes, loss of the insertional sequences causing fragmentation of nif genes and the loss of *nifHDKENB,* resulting in complete loss of nitrogen fixation capacity. It would be interesting to understand what environmental factors drive such evolutionary path toward loss of nitrogen fixation.

### Presence of Ancient nif and Ecological Context

Our analysis showed that an ancient *nif* (Group II) lineage, distinct from the canonical Group I *nif* found in most cyanobacteria, is distributed across 10 genera spanning multiple families. Although the presence of ancient *nif* in cyanobacteria has been previously suggested [12,58,69], our detailed analysis of these 10 genera identified 48 strains where both *nif* lineages coexist (Supplementary Table S11). These were categorized into three types: full coexistence of both *nif* types, complementary coexistence, and duplication of some genes. Metagenome profiling suggested that these strains primarily inhabit such as biofilms or microbial mats environments (Fig. 6). In such anaerobic habitats, active oxygen consumption via respiration by heterotrophic bacteria creates strictly anaerobic conditions [70].

Given that diazotrophic bacteria carrying the ancient *nif* are strict anaerobes, these cyanobacteria likely acquired their ancient *nif* via HGT from closely associated anaerobic bacteria within the microbial mats. The coexistence of both *nif* types suggests functional differentiation of these nif systems based on the environmental oxygen concentration: the canonical *nif* system is utilized when some oxic conditions by own oxygenic photosynthesis and the environment, while the ancient *nif* system operates under strict anaerobic conditions. Such differentiation may involve the master regulator CnfR. While CnfR was also conserved in strains carrying the ancient *nif* system, the conserved *cis*-acting elements recognized by CnfR in canonical *nif* systems (Tsujimoto et al., 2016)[48] were missing (Fig. 5). This suggests that the regulatory mechanism of CnfR in strains with the ancient *nif* genes is different from that of strains with the canonical *nif* genes to meet the different requirements of the ancient *nif* system.

### Gene Cluster Complexity

Based on the distribution of two types of the *nif* genes, we mapped the acquisition event of each *nif*-type within cyanobacterial lineage. The ubiquitous distribution of the canonical *nif* in cyanobacterial lineage except for Gloeobacter suggests that the canonical *nif* was acquired at the common ancestor that developed complex thylakoid membrane systems (Fig. 7, red arrow) [71]. In contrast, the ancient *nif* genes are found only in the four orders: Cyanobacteriales, Leptolyngbyales, Elainellales, and Phormidesmidales, suggesting the common ancestor of the four orders have acquired the ancient *nif* genes (Fig. 7, blue arrow). If this common ancestor already carried the canonical *nif*, the current diversity of the two *<u>nif</u>* types in the four orders implies dynamic evolutionary processes of coexistence, functional differentiation, or partial loss through genome streamlining.

**Figure 7.**
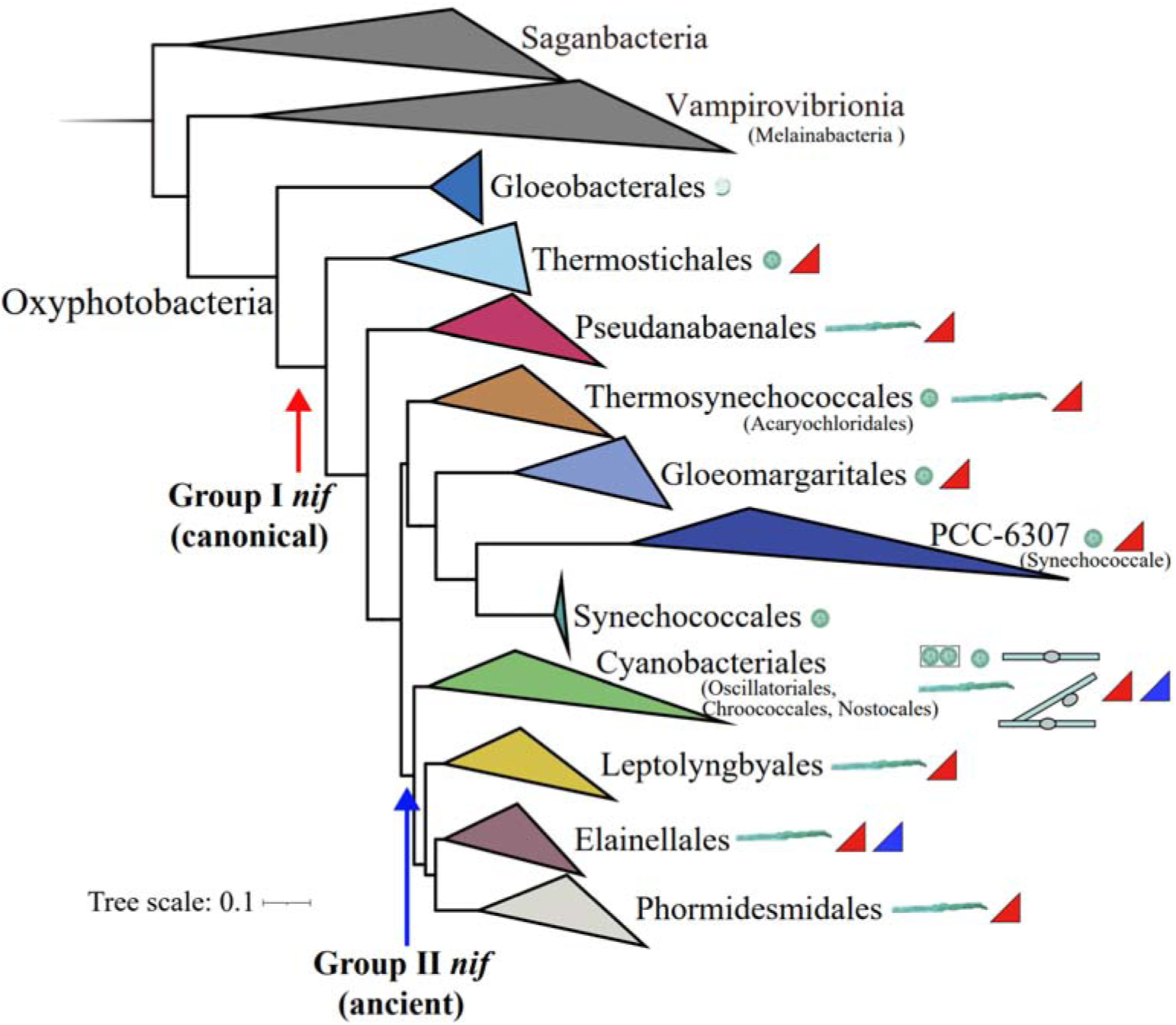
Probable evolutionary events gaining *nif* genes by HGT and distribution of Groups I and II *nif* genes in Cyanobacteria. A molecular phylogenetic tree of cyanobacteria at the order level, showing the morphology and *nif* types included, Group I (canonical, red triangles) and Group II (ancient, blue triangles). The canonical *nif* genes (Group I) were likely acquired at the base of the thylakoid-bearing lineages after divergence from the thylakoid-less *Gloeobacterales* (red arrow). The ancient *nif* genes (Group II) appear to be acquired at the base of later-branched lineages, with a potential common origin indicated by the blue arrow. GTDB classification and validly published nomenclature are used. Hypothetical orders are shown with validly published names provided in parentheses.

A comparison of gene cluster size revealed that canonical *nif* clusters contain significantly more genes (average 25) than ancient *nif* clusters (average 15, equivalent to *A. vinelandii*). This complexity may reflect the additional genetic element required to reconcile nitrogen fixation with oxygenic photosynthesis. These differences between them could provide critical clues for identifying the genes essential for control the Oxygen Paradox.

### Global FNF Patterns and Comparative Phylum Analysis

In the first curated 568-strain set, the FNF was 48.6% (50.5% excluding non-photosynthetic sister groups), which was consistent with the consensus that roughly half of cyanobacterial species are diazotrophs. However, when Nif-Finder applied to broader resources, the FNF decreased to 36.4% in the NCBI collection (1,365 genomes) and further to 19.4% in the GTDB collection (2,718 genomes). This suggests that the proportion of nitrogen-fixing cyanobacterial strains in nature may be much lower than previously estimated from cultured collections. Despite this, our estimation that Cyanobacteria is one of the phyla with the highest FNF across the bacterial domain, consistent with Koirala et al. (2021)[4], suggests that for photoautotrophic cyanobacteria, the ecological advantages of nitrogen fixation outweigh the metabolic costs of nitrogenase and the challenges of the Oxygen Paradox, maintaining a high FNF even through dynamic evolutionary processes of gains and losses.

### Future Perspectives

The *nifHDKENB* genes are considered the minimal unit for a functional nitrogenase (Dos Santos, 2012). However, there have been some cases in which even though carrying these genes no nitrogenase activity and no diazotrophic growth, as seen in *Vulcanococcus limneticus* LL. Successful integration of the *nif* genes into the preexisting host cyanobacterial metabolism requires at least solving the Oxygen Paradox. Engineering efforts to confer nitrogen fixation to *Synechocystis* sp. PCC 6803 by introduction of the relevant *nif* genes from diazotrophic cyanobacteria have achieved nitrogenase activity. However, no such transformants showed obvious nitrogen-fixing growth (Tsujimoto et al., 2018; Liu et al., 2018)[71,72]. This situation suggests that some additional genes beyond the *nifHDKENB* core genes are required for nitrogen fixing growth in oxygenic photosynthetic cells. The comprehensive distribution data and the comparative analysis of canonical versus ancient *nif* clusters provided in this study provide guide for identifying such missing essential genes.

## Materials and Methods

### Public cyanobacterial strains

3,620 and 2,818 cyanobacterial genomes were downloaded from the NCBI GenBank and GTDB [51] databases, respectively (accessed on June 30, 2022). Assembly quality was checked based on 105 single-copy marker genes using CheckM2 [73]. Following the MIMAG genomic standards [74], 2,199 genomes were retained based on the criteria of >90% completeness, <5% contamination, and <500 contigs. Taxonomic classification was performed using GTDB-tk v2.3 [75] with the R226 and R207 databases. These genomes were dereplicated with ANI 99% threshold using dRep [76] and further manually curated based on citation frequency or availability in major culture collections (Supplementary Fig. 2). Finally, 586 representative genomes were selected for *nif* gene search.

### Construction of Hidden Markov Model (HMM) profiles for nif gene search

A custom HMM profile was prepared to search for *nif* genes for highly divergent cyanobacterial lineages. First, amino acid sequences deduced from the *nifHDK* genes were downloaded from the nitrogen cycle gene database [77], including 14,758 NifH, 1,755 NifD, and 620 NifK sequences. Short sequences were removed, and remaining sequences were BLASTp [78] searched against UniRef100 [79]. Their taxonomic information was fetched using UniProt’s ID mapping service (https://www.uniprot.org/id-mapping/). Cyanobacterial Nif were clustered at 99% amino acid identity using CD-HIT v4.8 utility [80] to avoid oversampling of HMM profile. Their phylogenetic classification was inferred by ML method in IQ-TREE2 [81], with the domain superfamily of Nif protein as the outgroup. Finally, 98 NifH, 58 NifD, and 66 NifK sequences were selected (Supplementary Fig. 2). For each NifHDK sequence sets, profile HMMs were constructed using Hmmer3 suites [82]. As there are no NifENB sequences in the NCycDB database, their sequences were obtained from the Uniprot database. Full-length non-redundant cyanobacterial NifE, NifN, and NifB sequences were selected, and HMM profiles were constructed as described above (NifE: 61 proteins, NifN: 65 proteins, NifB: 54 proteins).

### nif/vnf gene search

Protein-coding genes were predicted for each genome using Prodigal [83] with default parameters. Hmmer3 suite [82] was used to search for NifHDKENB against 586 cyanobacterial proteomes with a prepared nif HMM profile (Supplementary Figure 2). All significant hits (<1E-5) were classified based on the BLASTp best hit to the SWISS-PROT sequence [84]. Phylogenetic inference was performed to assess the tree topology. To rescue the failure of gene prediction (e.g., fragmented *nif* in genomes of vegetative cells), six-frame translated BLASTx search [78] was also performed against 586 genomes, especially for Nostocaceae’s genomes. If fragmented *nif* were found, their genomic proximity (< 200kb), the order of N-terminus to C-terminus, and orientation of fragments were manually checked. VnfDKG were also searched to distinguish between V-type nitrogenase and Mo-type nitrogenase in the Nostocaceae family. Since V-type nitrogenases are monophyletic in Nostocaceae and not deeply diverged, single VnfDKG from *Anabaena variabilis* ATCC 29413 [85] was used as a query for BLASTx search against 168 Nostocaceae’s genomes. The presence or absence of *vnf* were determined by whether the Vnf monophyletic clade are observed in a Nif and Vnf mixed phylogenetic analysis.

### Phylogenetic inference

The evolutionary phylogeny of cyanobacterial strains was inferred using a concatenated set of 120 single copy bacterial marker genes defined in the GTDB [51]. Core gene extraction and multiple sequence alignment (MSA) construction were performed using the de_novo_wf method in GTDB-tk v2 [75] with the following options “--taxa_filter p Cyanobacteria --outgroup_taxon p Margulisbacteria -bacteria”. Phylogenetic tree reconstruction was performed using ML method in IQ-TREE v2.2.0.3 [81]. The best-fit substitution model was determined using ModelFinder [86] along with ultra-fast 1,000 bootstrap replication. The resulting circular phylogenetic tree was visualized on the iTOL v6 web service [87].

For Nif phylogenetic inference, Nif protein sequences were aligned using MAFFT with the “--maxiterate 100 --globalpair” options. Gapped sites were trimmed using trimAl (Capella-Gutiérrez et al., 2009) with the -gappyout option. Tree inference was subsequently conducted using IQ-TREE v2.2.0.3 with the options “-m MFP+MERGE -bb 1000 -alrt 1000”. For large datasets, FastTree v2 [88] was used instead, with the “-wag -gamma” options.

### Nif-Finder

On the 2D-Similarity plot of the negative log transformed *E*-value (−log_10_ E-value) versus protein length, NifHDKENB proteins formed a tight cluster, in contrast to the scattered distribution of NFL proteins (Fig. 1). This plot shows that each NifHDKENB protein is highly conserved within Cyanobacteria and possesses a distinct full-length primary sequence compared with NFLs. Consequently, cyanobacterial Nif and NFL proteins can be accurately classified on the 2D-Similarity plot. We implemented a simple Python script to rapidly identify *NifHDKENB* from cyanobacterial protein datasets. The program uses user-provided protein sequences as queries and predicts Nif/Vnf through two steps.

1. HMMER3 search using custom Nif profiles: User-provided protein sequences (FASTA format) are screened against custom cyanobacterial NifHDKENB HMM profiles using hmmscan. For each significant hit, the −log_10_ E-value and the hit protein length are extracted from hmmscan result.
2. Distance-based annotation: The −log_10_ E-value and protein length of each query are compared against those of the Nif and NFL proteins from the 586 reference dataset determined in this study (Fig. 1). The annotation of the query protein is determined by calculating the straight distance (*d*) between the query and reference proteins in a two-dimensional space. The distance *d_i_,_r_* between query *i* and reference protein *r* is defined as:

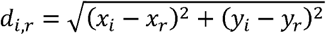

Here, *x* means the protein length and *y* means the –log_10_ E-value. The annotation of the closest reference is assigned to the query based on the minimum straight distance (Euclidean distance) without any additional transformation or normalization.

Here, genomes in which all NifHDKENB components are assigned as full-length are classified as potential diazotrophs.

### Identification of nif-clusters

First, genomic positions of *nifHDK* were identified using Nif-Finder. Then, thirty ORFs flanking *nifHDK* locus were extracted from the GFF files annotated with Bakta [89]. Then, synteny among extracted ORFs was visually examined using Clinker [90] with default settings. ORFs that showed no similarity to other *nif*-clusters or to surrounding ORFs were trimmed.

## Data Availability

Nif-Finder is implemented in Python and runs on Linux, macOS, or WSL environments. Nif screening of 586 strains with manual curation took two months, whereas Nif-Finder completed the screening of the 143,619 representative bacteria in GTDB R226 in a single day (when run in 20 parallel runs). The Nif-Finder program and Cyanobacterial nif HMM profiles is freely available at https://github.com/kazumaxneo/Nif_finder under the GPL v3 license, along with 285 cyanobacterial Nif HMM profile sets and other diazotrophic bacterial group I Nif profiles.

## Acknowledgments

We thank Kunio Ihara for use of computational resources in Center for Gene Research in Nagoya University. We also thank Mari Banba, Haruki Yamamoto, Takafumi Yamashino and all members of the Laboratory of Molecular and Functional Genomics in Nagoya University for engaging in discussion.

## Author contributions

### Disclosures

The authors have no conflicts of interest to declare.

