## Supplemental Figures for "Accurate prediction of nitrogen fixation in cyanobacteria reveals the dynamic evolution driving high retention rate with mosaic distribution"

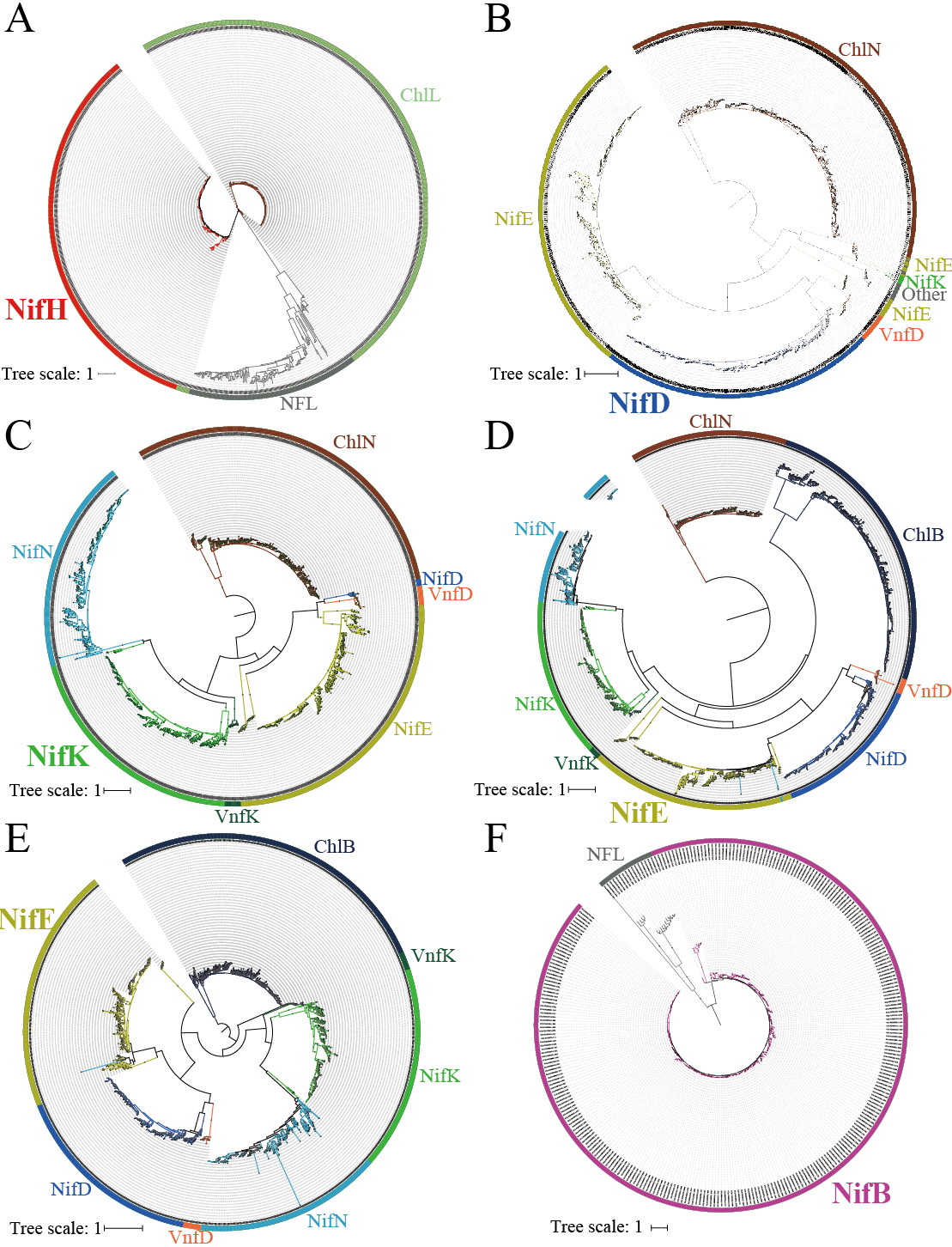

**Supplementary Figure S1.** Confirmation of the Nif orthologs in the Nif superfamily proteins identified by homology searches against 586 cyanobacterial strains by phylogenetic analysis using cyanobacterial HMM profiles. Each circular plot shows significant hits (E < 1 × 10⁻^5^) obtained using HMM profiles of NifH (**A**), NifD (**B**), NifK (**C**), NifE (**D**), NifN (**E**), and NifB (**F**). The color of the outermost ring indicates the single best hit in the SWISS-PROT database and corresponds to the colors used in the Nif 2D Similarity plot in Figure 1.

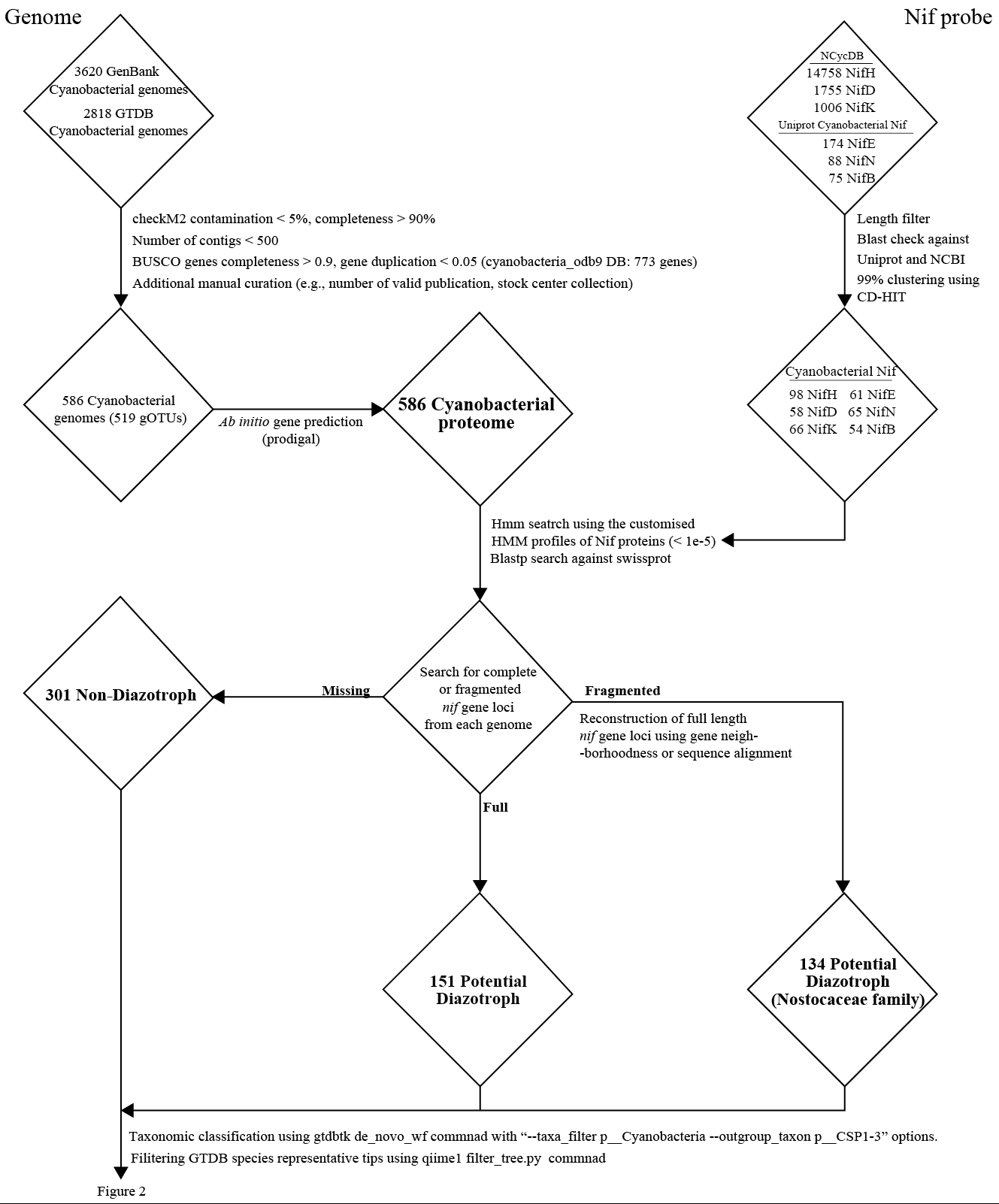
**Supplementary Figure S2.** Flowchart for the *nifHDKENB* identification from 586 cyanobacterial strains.

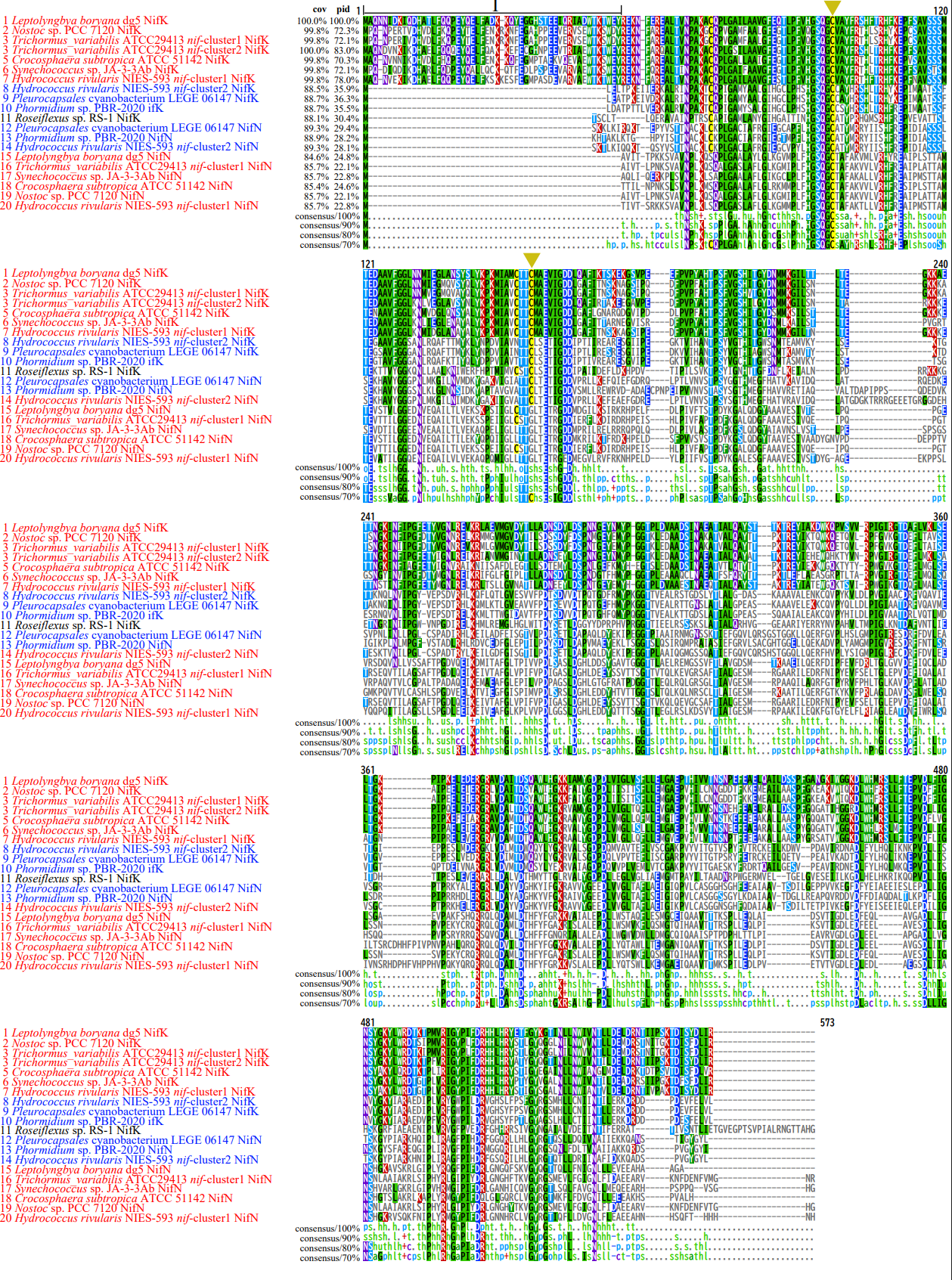

**Supplementary Figure S3.** Multiple sequence alignment of cyanobacterial NifK and NifN proteins. Groups I and II Nif proteins are shown in red and blue, respectively. Group I NifK contains a long N-terminal extension ([Zuviría et al, 2025](https://pmc.ncbi.nlm.nih.gov/articles/PMC12425478/" \l "bib21)). Multiple sequence alignment was performed using MAFFT (Katoh and Standley., 2013) with the “--globalpair --maxiterate 100 --thread 8” options.

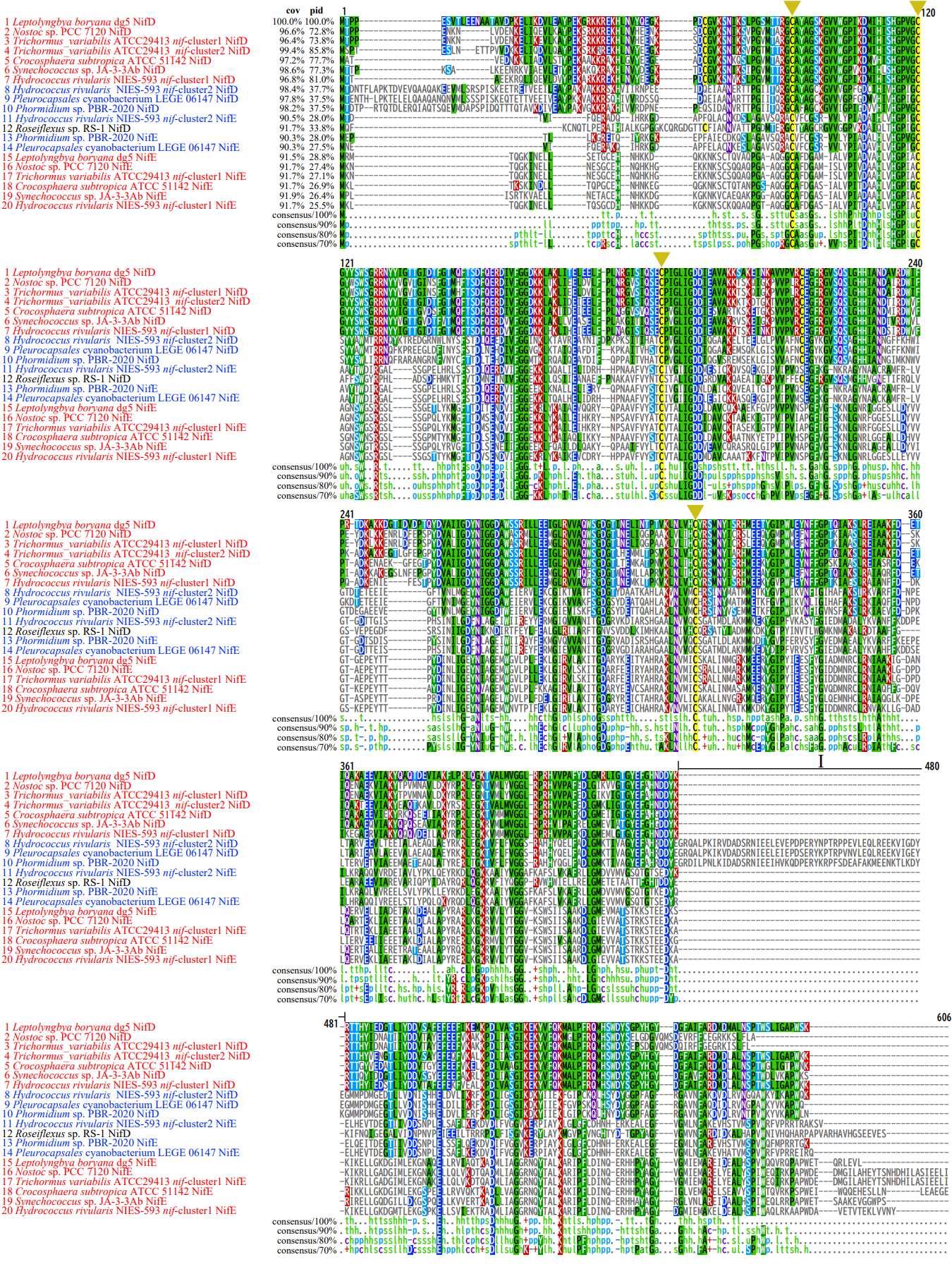

**Supplementary Figure S4.** Multiple sequence alignment of cyanobacterial NifD and NifE proteins. Groups I and II Nif proteins are shown in red and blue, respectively. Group II NifD contains a long insertion (I) ([Zuviría et al, 2025](https://pmc.ncbi.nlm.nih.gov/articles/PMC12425478/" \l "bib21)). Multiple sequence alignment was performed using MAFFT (Katoh and Standley., 2013) with the “--globalpair --maxiterate 100 --thread 8” options.

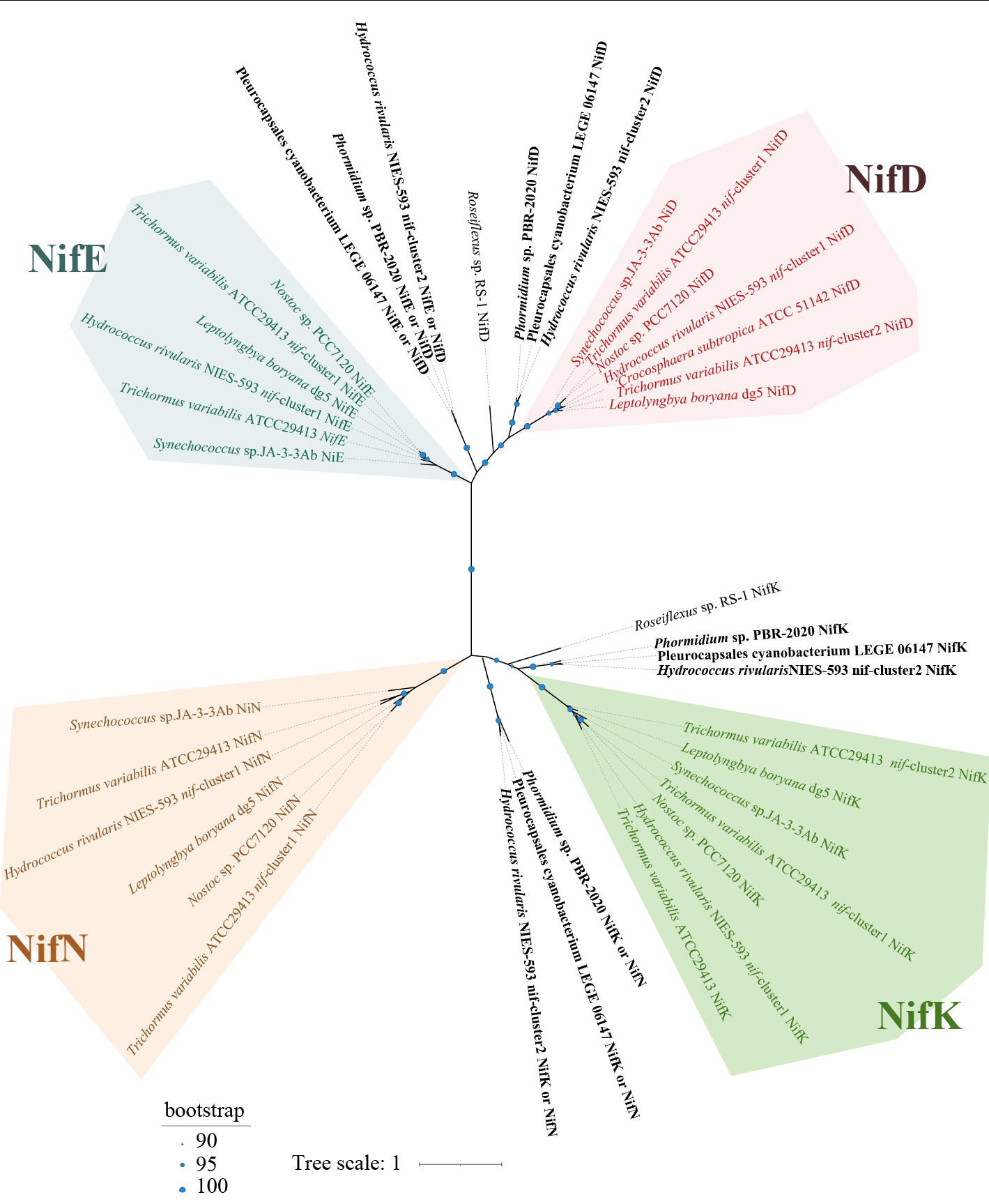

**Supplementary Figure S5.** Unrooted phylogenetic tree of the canonical and ancestral NifDKEN proteins. As the representatives of canonical Nif (Group I), six sets of NifDKEN proteins from *Trichormus variabilis* ATCC 29413 (two copies), *Nostoc* sp. PCC 7120, *Leptolyngbya boryana* dg5, *Synechococcus* sp. JA-3-3b, and Group I *nif* of *Hydrococcus rivularis* NIES-593 were used (colored). As the representatives of ancestral Nif (Group II), three sets of NifDKEN proteins from *Phormidium* sp. PBR-2020, Pleurocapsales cyanobacterium LEGE 06147, and Group II *nif* of *Hydrococcus rivularis* NIES-593 were used (black). Canonical NifDKEN proteins are highlighted in color as in Figure 4. Tree inference was performed using the maximum likelihood method with 1,000 ultrafast bootstrap replicates in IQ-TREE 3 (Wang et al., 2025), with the best-fit substitution models LG+I+G4 as inferred by ModelFinder-Plus (Kalyaanamoorthy et al., 2017).

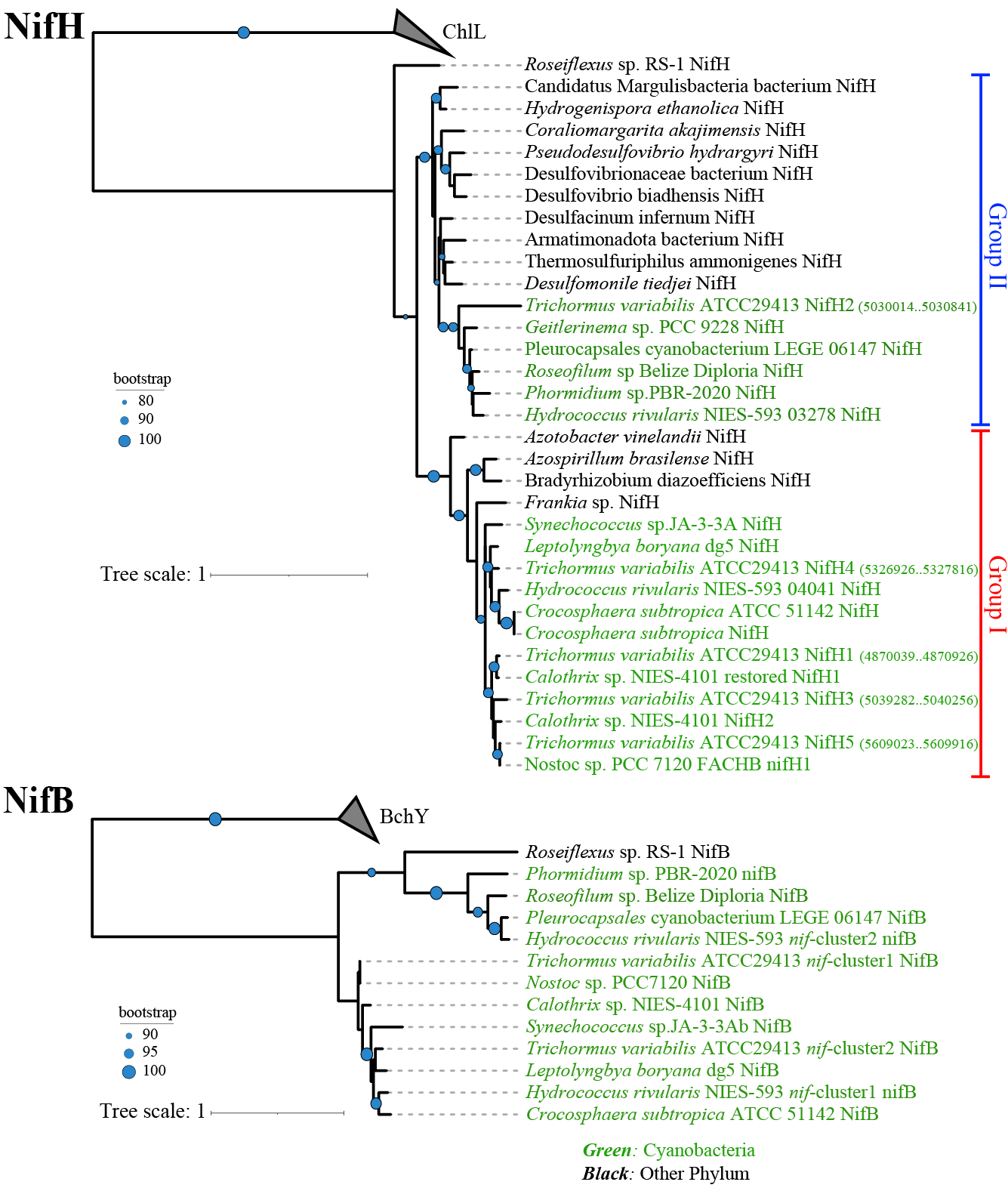

**Supplementary Figure S6.** Rectangular phylogenetic trees of canonical and ancestral Nif proteins. NifH (upper) and NifB (lower) trees are shown. In each tree, cyanobacterial Nif proteins are highlighted in green. Phylogenetic inference was performed using the maximum likelihood method with 1,000 ultrafast bootstrap replicates implemented in IQ-TREE 3 (wang et al., 2025), using the best-fit substitution models LG+I+R3 for NifH and LG+G4 for NifB as determined by ModelFinder-Plus (Kalyaanamoorthy et al., 2017).

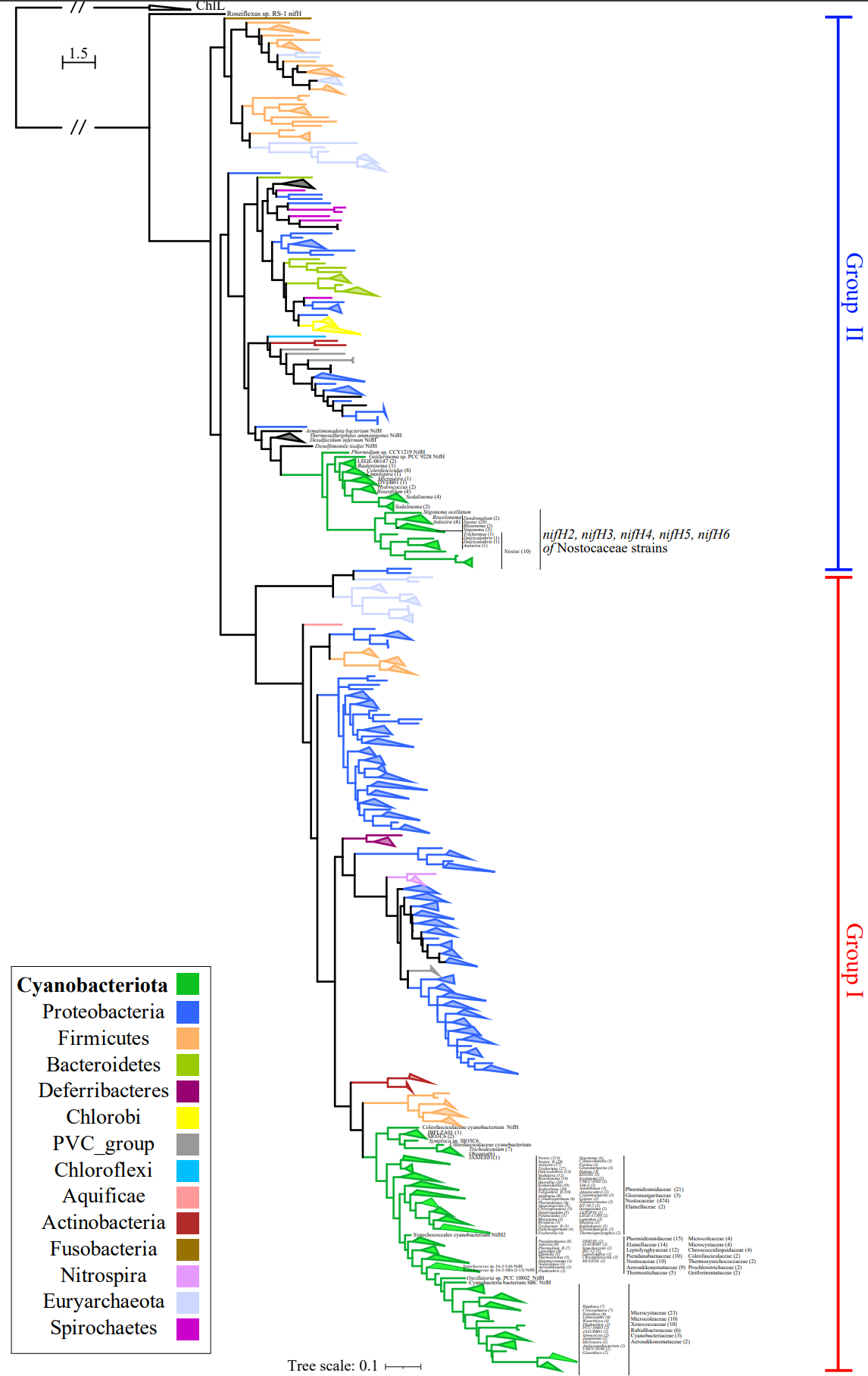

**Supplementary Figure S7.** Rectangular phylogenetic trees of NifH proteins in Groups I and II. Colored nodes indicate phylum-level taxonomy. Cyanobacterial NifH proteins (green) are classified into Groups I (canonical) and II (ancestral) clades. Phylogenetic inference was performed using the approximately maximum-likelihood method implemented in FastTree (Price et al., 2009), with the WAG substitution model and gamma-distributed rate heterogeneity among sites. Branches with bootstrap resampling support ≤ 0.1 were collapsed on the iTOL v.6 web-server.

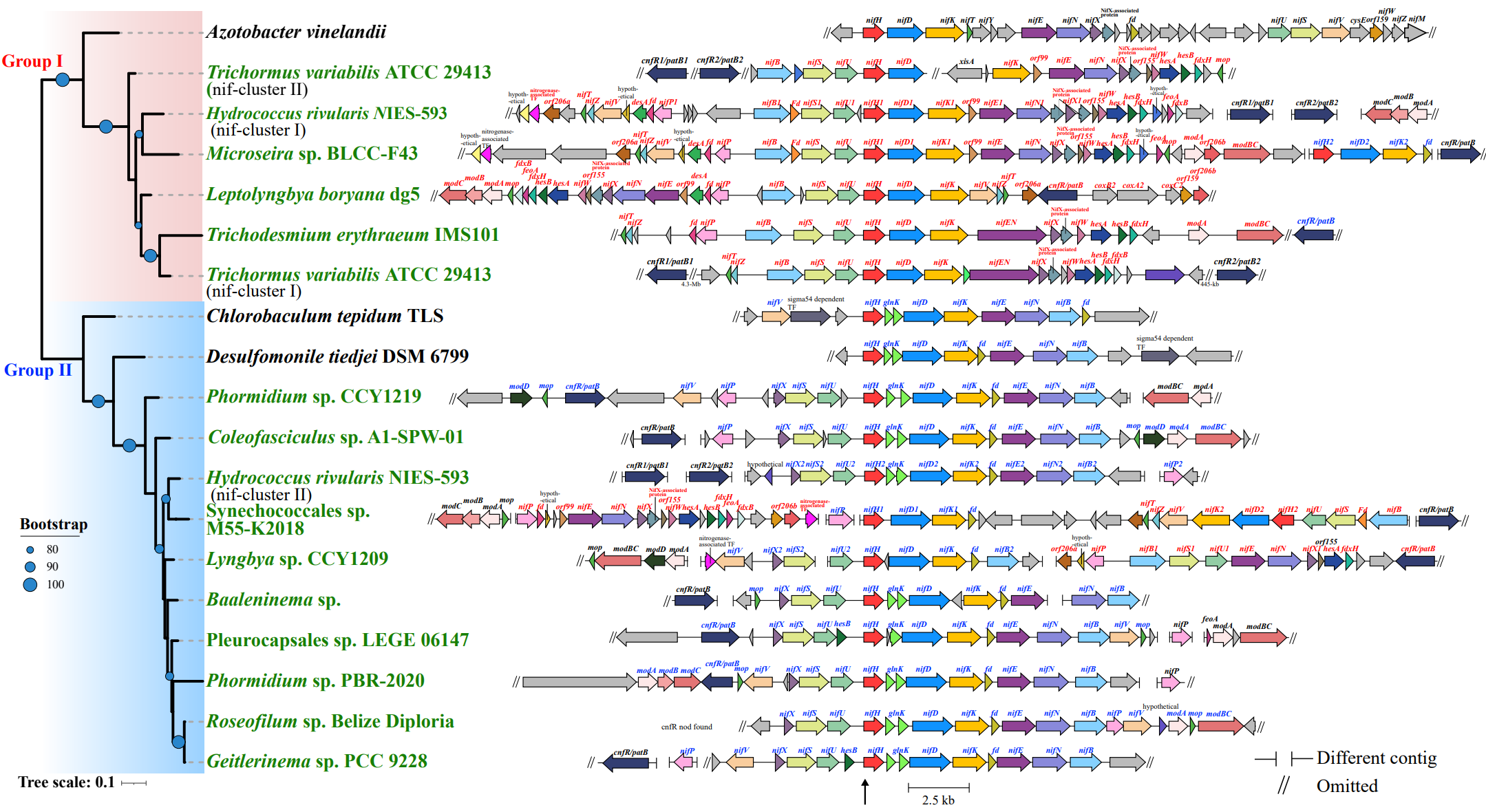

**Supplementary Figure S8.** Comparison of gene arrangement of the *nif* gene clusters in Groups I and II. ORFs are shown as thick horizontal arrows, with identical colors indicating orthologous genes. Gene names of Groups I and II are highlighted in red and blue, respectively. *Synechococcales* sp. M55-K2018 and *Microseira* sp. BLCC-F43 encodes both Group I (canonical) *nifHDKENB* and Group II (ancestral) *nifHDK*. *Lyngbya* sp. CCY1209 encodes Group I *nifENB* and Group II *nifHDKB*.

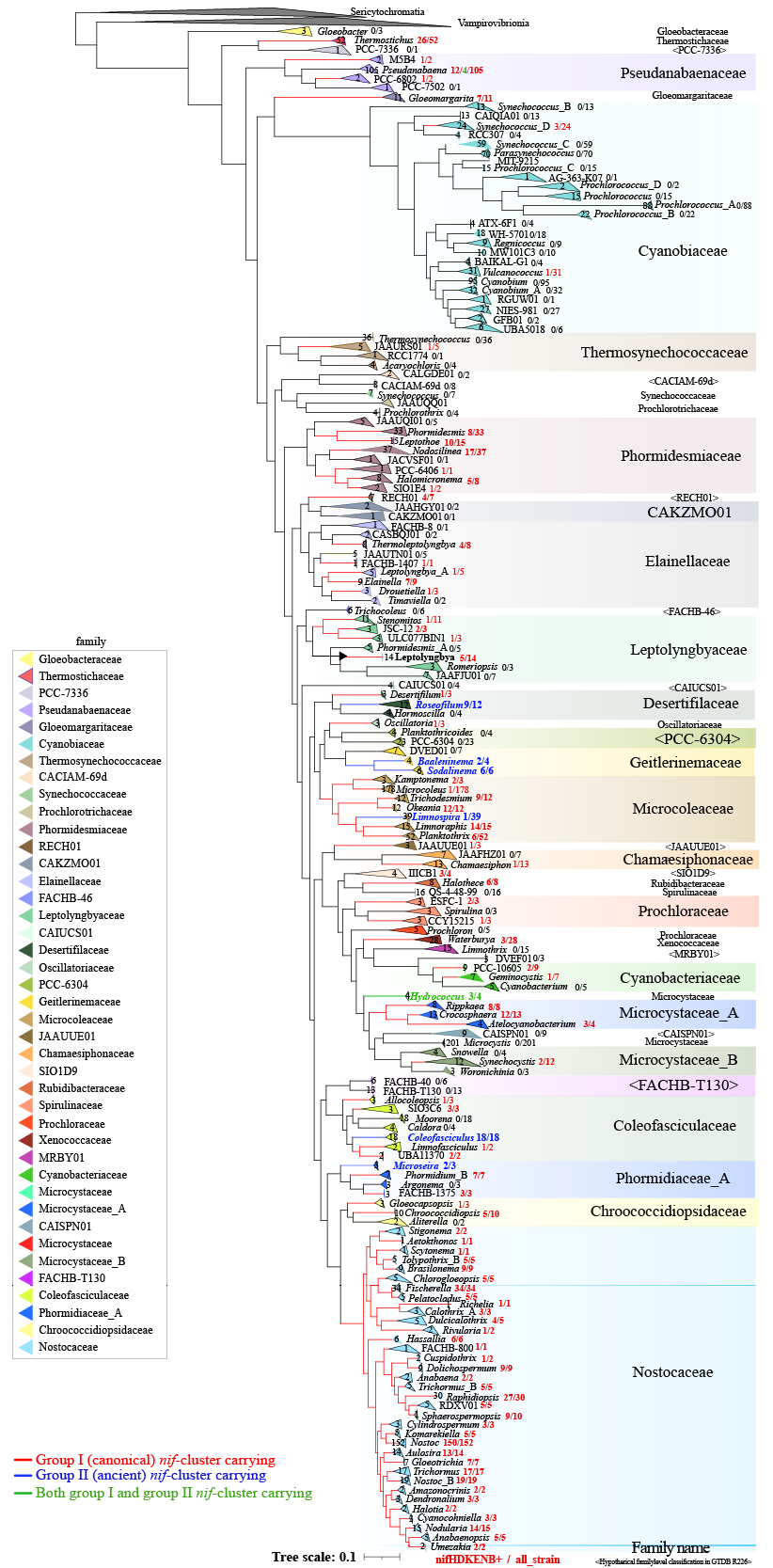

**Supplementary Figure S9.** Cyanobacterial phylogenetic tree at genus level showing the distribution of canonical (Group I) and ancestral (Group II) *nif* carrying lineages. A total of 2,716 representative strains from GTDB R226 were analyzed, excluding GTDB hypothetical genera. Phylogenetic inference was performed using the approximately ML method implemented in FastTree (Price et al., 2009), with the WAG substitution model and gamma-distributed rate heterogeneity among sites. The analysis was based on approximately 5,000 highly conserved, gap-free sites derived from the 120 core genes defined by GTDB (Parks et al., 2018). Family-level taxonomic classifications defined in R226 are highlighted with background colors. The red, blue and green digits indicate the number of strains carrying canonical (Group I), ancestral (Group II) and both *nif* genes, respectively; relative to the total number of strains in each genus identified by Nif-Finder. The full-size figure is available from the Nif-Finder repository.

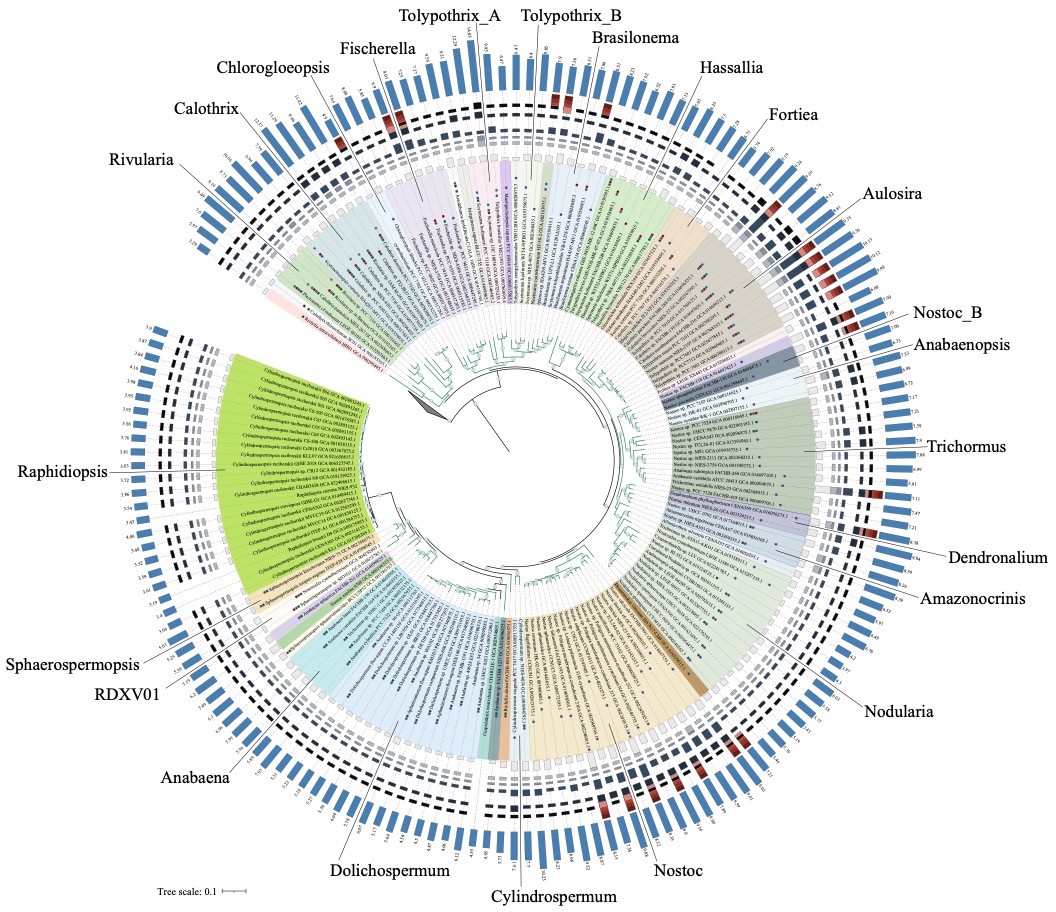

**Supplementary Figure S10.** Comparison of genome size and *nif*/*vnf* copy numbers among Nostocaceae clades. The ML tree corresponding to the Nostocaceae clade was extracted from the ML tree in Figure 2A. Genome sizes of each strain are shown in the outermost blue colored box plot. Copy numbers of *nif* and *vnf* from 167 Nostocaceae strains are shown in the outer box plots (gray, MoFe-type *nifHDKENB*; orange, VFe-type *vnfDKG*).
